# Role of endopeptidases in peptidoglycan synthesis mediated by alternative cross-linking enzymes in *Escherichia coli*

**DOI:** 10.1101/2021.02.12.430937

**Authors:** Henri Voedts, Delphine Dorchêne, Adam Lodge, Waldemar Vollmer, Michel Arthur, Jean-Emmanuel Hugonnet

## Abstract

Bacteria resist to the turgor pressure of the cytoplasm through a net-like macromolecule, the peptidoglycan, made of glycan strands connected via peptides cross-linked by penicillin-binding proteins (PBPs). We recently reported the emergence of β-lactam resistance resulting from a bypass of PBPs by the YcbB L,D-transpeptidase (LdtD), which form chemically distinct 3→3 cross-links compared to 4→3 formed by PBPs. Here we show that peptidoglycan expansion requires controlled hydrolysis of cross-links and identify amongst eight endopeptidase paralogues the minimum enzyme complements essential for bacterial growth with 4→3 (MepM) and 3→3 (MepM and MepK) cross-links. Purified Mep endopeptidases unexpectedly displayed a 4→3 and 3→3 dual specificity implying recognition of a common motif in the two cross-link types. Uncoupling of the polymerization of glycan chains from the 4→3 cross-linking reaction was found to facilitate the bypass of PBPs by YcbB. These results illustrate the plasticity of the peptidoglycan polymerization machinery in response to the selective pressure of β-lactams.

## INTRODUCTION

Peptidoglycan (PG) is an essential macromolecule that surrounds the bacterial cell providing resistance to the osmotic pressure of the cytoplasm and determining cell shape (Turner et al., 2014). PG is assembled from a disaccharide-peptide subunit consisting of *N*-acetylglucosamine (GlcNAc) and *N*-acetylmuramic acid (MurNAc) substituted by a stem pentapeptide (L-Ala^1^-γ-D-Glu^2^-DAP^3^-D-Ala^4^-D-Ala^5^ in which DAP is diaminopimelic acid) (Fig. 1A). The subunit is assembled by glycosyltransferases that polymerize glycan strands and transpeptidases that form amide bonds between stem peptides carried by adjacent glycan strands. *Escherichia coli* relies on two types of transpeptidases for the latter reaction (Magnet et al., 2008). The D,D-transpeptidases, also referred to as penicillin-binding proteins (PBPs), form the most abundant cross-links, which connect the fourth residue (D-Ala^4^) of an acyl donor to the third residue (DAP^3^) of an acyl acceptor (4→3 cross-link) (Fig. 1A). The L,D-transpeptidases (LDTs) form 3→3 cross-links that connect two DAP residues (Fig. 1B). The D-Ala at the 5^th^ and 4^th^ positions of stem peptides that do not participate in cross-link formation as donors are fully and partially trimmed by carboxypeptidases of the D,D and L,D specificities, respectively (Fig. 1C). PBPs and LDTs are structurally unrelated, rely on different catalytic nucleophiles (Ser *versus* Cys, respectively), and use different acyl donor stems (pentapeptide *versus* tetrapeptide, respectively) (Mainardi et al., 2008; Sauvage et al., 2008). PBPs and LDTs also differ by their inhibition profiles since PBPs are potentially inhibited by all classes of β-lactams (including penams, cephems, monobactams, and carbapenems) whereas LDTs are effectively inhibited only by carbapenems (Mainardi et al., 2005). LDTs are fully dispensable for growth of *E. coli*, at least in laboratory conditions, and form a minority of the cross-links during exponential growth (6% of the total cross-links) (Glauner et al., 1988; Sanders and Pavelka, 2013). The proportion of 3→3 cross-links is higher in the stationary phase (Pisabarro et al., 1985) and in cells experiencing outer membrane assembly stress (Morè et al., 2019). Bypass of PBPs by LDTs leads to high-level resistance to β-lactams of the penam (such as ampicillin), cephem (ceftriaxone), and monobactam (aztreonam) classes in engineered *E. coli* strains that overproduce the YcbB L,D-transpeptidase, also referred to as LdtD, and the guanosine penta- and tetra-phosphate [(p)ppGpp] alarmones (Hugonnet et al., 2016). PG of such strains grown in the presence of β-lactams exclusively contains 3→3 cross-links, indicating that the D,D-transpeptidase activity of PBPs is fully replaced by the L,D-transpeptidase activity of LDTs.

**Figure 1.**
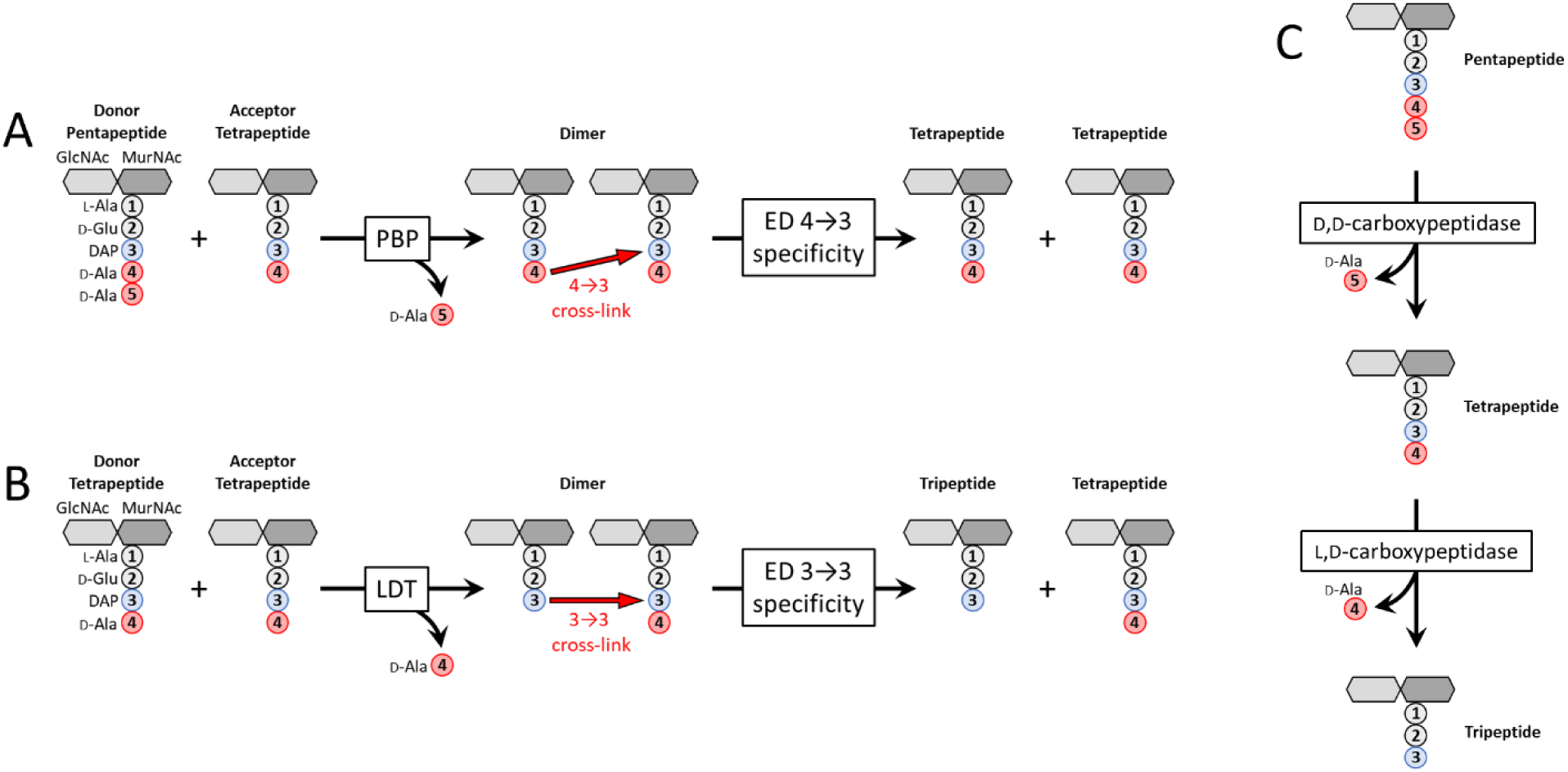
Metabolism of PG cross-links and maturation of free stem peptides. Formation and hydrolysis of (**A**) 4→3 and (**B**) 3→3 cross-links. The disaccharide-pentapeptide unit is assembled from *N-* acetylglucosamine (GlcNAc), *N*-acetylmuramic acid (MurNAc), and five amino acids including *meso-* diaminopimelic acid (DAP), which is linked via its L (S) center to the γ-carboxyl group of D-Glu. (**C**) Hydrolysis of the D-Ala^4^-D-Ala^5^ and DAP^3^-D-Ala^4^ peptide bonds by carboxypeptidases of D,D and L,D specificities, respectively.

**Figure S1.**
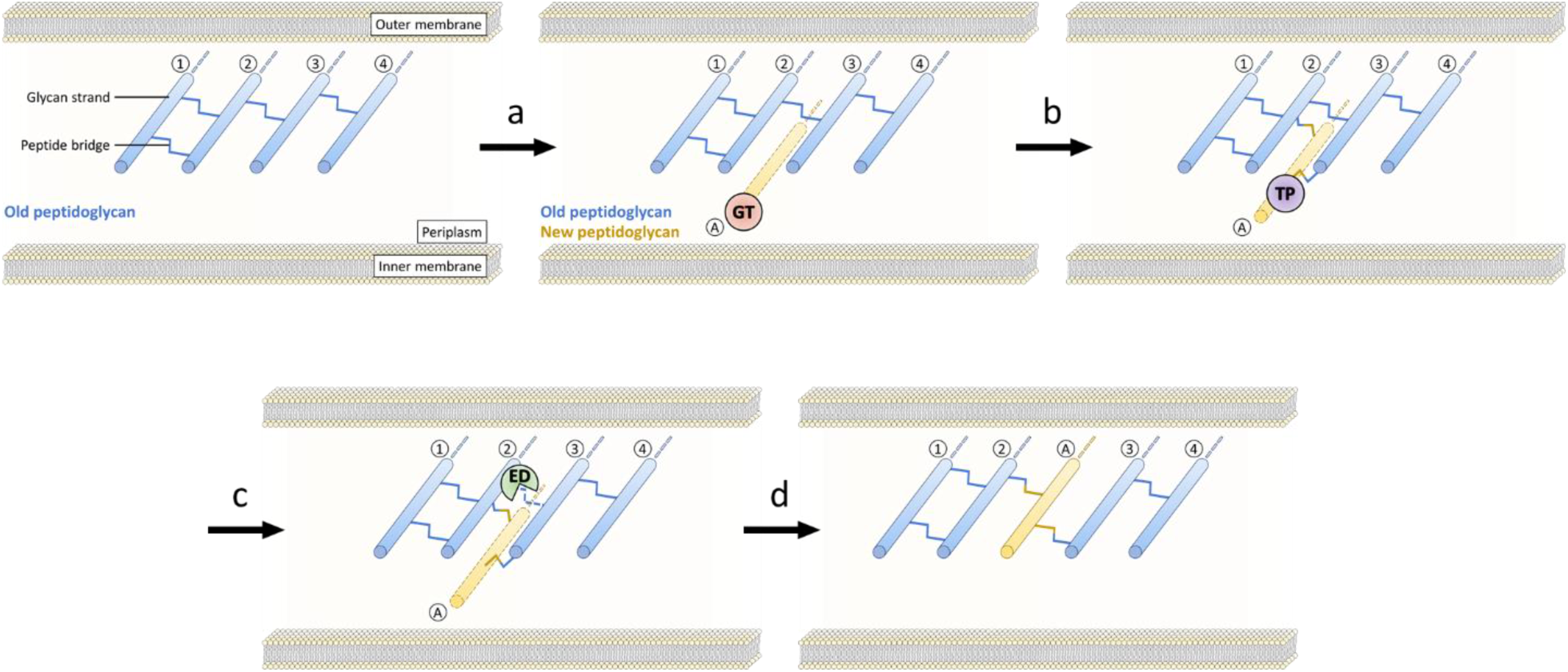
Insertion of PG subunits into the growing PG network. According to this model, one glycan strand (A) is polymerized by glycosyltransferases (GTs; step a) and attached to the pre-existing polymer (strands 2 and 3) by transpeptidases (TPs; step b). Hydrolysis of the cross-links connecting strands 2 and 3 by endopeptidases (EDs; step c) results in the expansion of the PG layer (step d). Of note, this model, which applies to the synthesis of the lateral wall, accounts for incorporation of new subunits sheltered from the cytoplasm osmotic pressure.

Expansion of PG is thought to require highly regulated hydrolytic activities that spatially control the insertion of new subunits into the growing network of cross-linked glycan strands (Singh et al., 2012; Vollmer, 2012). Due to the covalent net-like structure of PG, endopeptidases are predicted to be required for insertion of new subunits leading to the expansion of the PG layer (supplementary Fig. S1) (Höltje and Heidrich, 2001). The *E. coli* genome encodes eight endopeptidase paralogues that belong to five enzyme families (Chodisetti and Reddy, 2019; Pazos and Peters, 2019; Singh et al., 2012). PBP4, PBP7, and AmpH belong to the acyl-serine transferase superfamily, which also comprises D,D-transpeptidases and D,D-carboxypeptidases. Members of this superfamily are inhibited by β-lactam antibiotics. The NIpC/P60 cysteine peptidase family comprises two paralogues (MepH and MepS). Metallo-enzymes are represented by three enzyme families, LAS metallopeptidases, lysostaphin/M23 peptidases, and M15 peptidases, each contributing one paralogue (MepA, MepM, and MepK, respectively). The specificity of these eight paralogues as endopeptidases or carboxypeptidases (Fig. 1) has been explored by using sacculi or purified PG fragments as substrates (Chodisetti and Reddy, 2019; Engel et al., 1992; Gonzalez-Leiza et al., 2011; Keck and Schwarz, 1979; Korat et al., 1991; Romeis and Holtje, 1994; Singh et al., 2012). PBP4, PBP7, AmpH, MepH, and MepS hydrolyze 4→3 cross-links but PG dimers containing 3→3 cross-links were not tested. In contrast, MepA, MepM, and MepK were fully characterized revealing that MepA hydrolyzes both 4→3 and 3→3 cross-links, MepM is specific to 4→3 cross-links, and MepK displays a marked preference for 3→3 cross-links. The endopeptidases of *E. coli* are redundant and their essential roles can only be revealed by introducing multiple chromosomal deletions. One study unambiguously showed that hydrolysis of 4→3 cross-links by endopeptidases is essential as the triple deletion of genes encoding MepH, MepM, and MepS was not compatible with growth of *E. coli* in laboratory conditions (Singh et al., 2012; comment by Vollmer, 2012). Several endopeptidases interact genetically and physically with the outer membrane anchored adaptor protein NlpI supporting overlapping functions during the cell cycle (Banzhaf et al., 2020).

The dual capacity of *E. coli* to use transpeptidases of the D,D and L,D specificities raises the possibility that polymerization of PG containing 4→3 or 3→3 cross-links involves two overlapping sets of endopeptidases. To address this question, we used an *E. coli* strain that conditionally and exclusively relies on the formation of 3→3 cross-links for growth in the presence of ampicillin or ceftriaxone (Hugonnet et al., 2016). By introducing serial deletions of endopeptidase genes, we showed that the 4→3 and 3→3 modes of PG polymerization both require hydrolysis of cross-links. We identified distinct sets of endopeptidases that are essential for growth involving the two modes of PG cross-linking. Strikingly, impaired digestion of nascent glycan strands by a lytic transglycosylase was found to favor PG polymerization mediated by LDTs. These results highlight the functional plasticity of PG polymerization complexes to accommodate various PG cross-linking enzymes and hydrolases.

## RESULTS

### MepM is essential for β-lactam resistance mediated by the YcbB L,D-transpeptidase

The role of endopeptidases was assessed in *E. coli* BW25113 *ΔrelA pKT2(ycbB) pKT8(relA’)*, BW25113(*ycbB*, *relA’)* in short, which enables controlling the relative contribution of formation of 4→3 and 3→3 cross-links to PG polymerization (Hugonnet et al., 2016). In this strain, the *ycbB* L,D-transpeptidase gene carried by plasmid pKT2 is expressed under the control of an IPTG-inducible promoter. Plasmid pKT8 carries an L-arabinose-inducible copy of the *relA’* gene encoding a truncated version of RelA (residues 1 to 455), which synthesizes the (p)ppGpp alarmone in an unregulated manner due to the absence of the C-terminal ribosome binding module (Schreiber et al., 1991). In the presence of both inducers, production of YcbB and RelA’ is sufficient for full bypass of the D,D-transpeptidase activity of PBPs by the L,D-transpeptidase activity of YcbB (Fig. 2A). This enables bacterial growth in the presence of ampicillin or ceftriaxone since these drugs do not inhibit the YcbB L,D-transpeptidase. Testing for the inducible expression of β-lactam resistance in BW25113(*ycbB*, *relA’)* therefore provides a means to identify genes that are essential for growth when PG cross-linking is exclusively mediated by the YcbB L,D-transpeptidase. We used this phenotypic assay to assess the individual role of seven of the eight endopeptidases of *E. coli* following single-gene deletions in the BW25113(*ycbB*, *relA’)* strain (Fig. 2B). The remaining endopeptidase MepK could not be tested by this approach since deletion of the corresponding gene was not compatible with the presence of plasmid pKT2(*ycbB*) (see below). Mutants with deletion of *mepA, mepH, dacB, pbpG*, or *ampH* were resistant to ceftriaxone. Deletion of *mepS* decreased plating efficiency in the presence of the drug. Deletion of *mepM* abolished expression of β-lactam resistance. These results are surprising since MepM and MepS were not reported to hydrolyze 3→3 cross-links (Chodisetti and Reddy, 2019; Singh et al., 2012).

**Figure 2.**
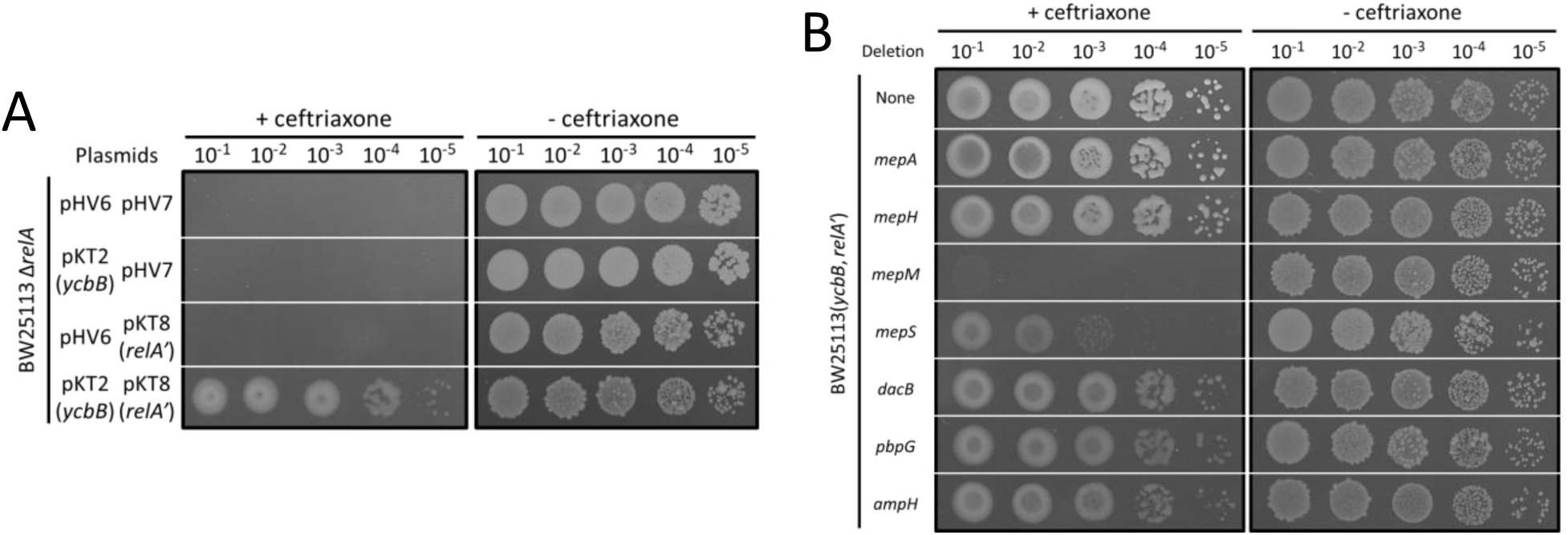
MepM is essential for YcbB-mediated β-lactam resistance. Growth was tested in the presence of ceftriaxone at 8 μg/ml (+ ceftriaxone) or in the absence of the drug (-ceftriaxone) on BHI agar plates supplemented with 40 μM IPTG and 1% L-arabinose for induction of *ycbB* and *relA’*, respectively. (**A**) BW25113 *ΔrelA* derivatives harboring plasmids pKT2(*ycbB*), pKT8(*relA’*) and the vectors pHV6 and pHV7 used to construct these plasmids, respectively. Expression of β-lactam resistance requires induction of both *ycbB* and *relA’*. (**B**) BW25113(*ycbB*, *relA’)* and its derivatives obtained by individual deletion of endopeptidase genes. BW25113(*ycbB*, *relA’)* is an abbreviated name for BW25113 *ΔrelA* pKT2(*ycbB*) pKT8(*relA’*).

### Production of YcbB is lethal in the absence of MepK

The gene encoding MepK, an endopeptidase with the dual 4→3 and 3→3 specificities was readily deleted from the chromosome of BW25113 *ΔrelA*. The resulting strain, BW25113 *ΔrelA ΔmepK* was transformed with pKT2(*ycbB*), pKT8(*relA’*), or both plasmids in combination (co-transformation). Tetracycline and chloramphenicol were used to select transformants that acquired pKT2(*ycbB*) and pKT8(*relA’*), respectively. Plasmid pKT8(*relA’*) was readily introduced into BW25113 Δ*relA* Δ*mepK* by transformation (10^8^ transformants per μg of DNA). Plasmid pKT2(*ycbB*) alone or in combination with pKT8(*relA’*) could not be introduced into BW25113 *ΔrelA ΔmepK* (< 5 transformants per μg of DNA). The same plating efficacies were observed in selective media containing IPTG, L-arabinose, or both inducers, in addition to tetracycline and chloramphenicol. These results show that production of the YcbB L,D-transpeptidase is lethal in the absence of MepK, in agreement with a recent report (Chodisetti and Reddy, 2019). Thus, cleavage of 3→3 cross-links by MepK is essential for bacterial growth when the proportion of 3→3 cross-links is increased in the presence of a plasmid copy of *ycbB*. Quantitatively, the basal level of *ycbB* expression in the absence of IPTG was sufficient for the lethal phenotype associated with the *mepK* deletion. Under non-inducing conditions the relative proportion of 4→3 and 3→3 cross-links in the PG extracted from exponential phase cultures of *BW25113(ycbB, relA’)* was in the order of 60% and 40%, respectively (data not shown). Thus, the cleavage of 3→3 cross-links by MepK was essential even if these cross-links co-existed with 4→3 cross-links formed by the PBPs.

### The hydrolytic activity of MepM is essential for β-lactam resistance

Deletion of the *mepM* gene abolished YcbB-mediated β-lactam resistance (above, Fig. 2B) even though this endopeptidase was not reported to cleave 3→3 cross-links (Chodisetti and Reddy, 2019; Singh et al., 2012). We therefore considered the possibility that the essential role of *mepM* in resistance could involve an as yet unknown function in addition to its 4→3-endopeptidase activity. MepM (440 residues) comprises a LytM (lysostaphin/M23 peptidase) domain and a LysM PG-binding domain (Pfam: P0AFS9). Complementation analysis of the *mepM* deletion in BW25113(*ycbB*, *relA’)* was performed with plasmids encoding (i) MepM, (ii) MepM H^393^A harboring an Ala residue at position 393 in place of an essential catalytic His residue conserved in members of the M23 peptidase family, and (iii) MepM ΔLytM lacking the C-terminal endopeptidase catalytic domain (Fig. 3). Expression of β-lactam resistance by BW25113(*ycbB*, *relA’) ΔmepM* was only restored by the plasmid harboring an intact copy of the *mepM* gene. Thus, the complementation analysis led to the conclusion that the endopeptidase activity of MepM is essential for YcbB-mediated resistance to β-lactams.

**Figure 3.**
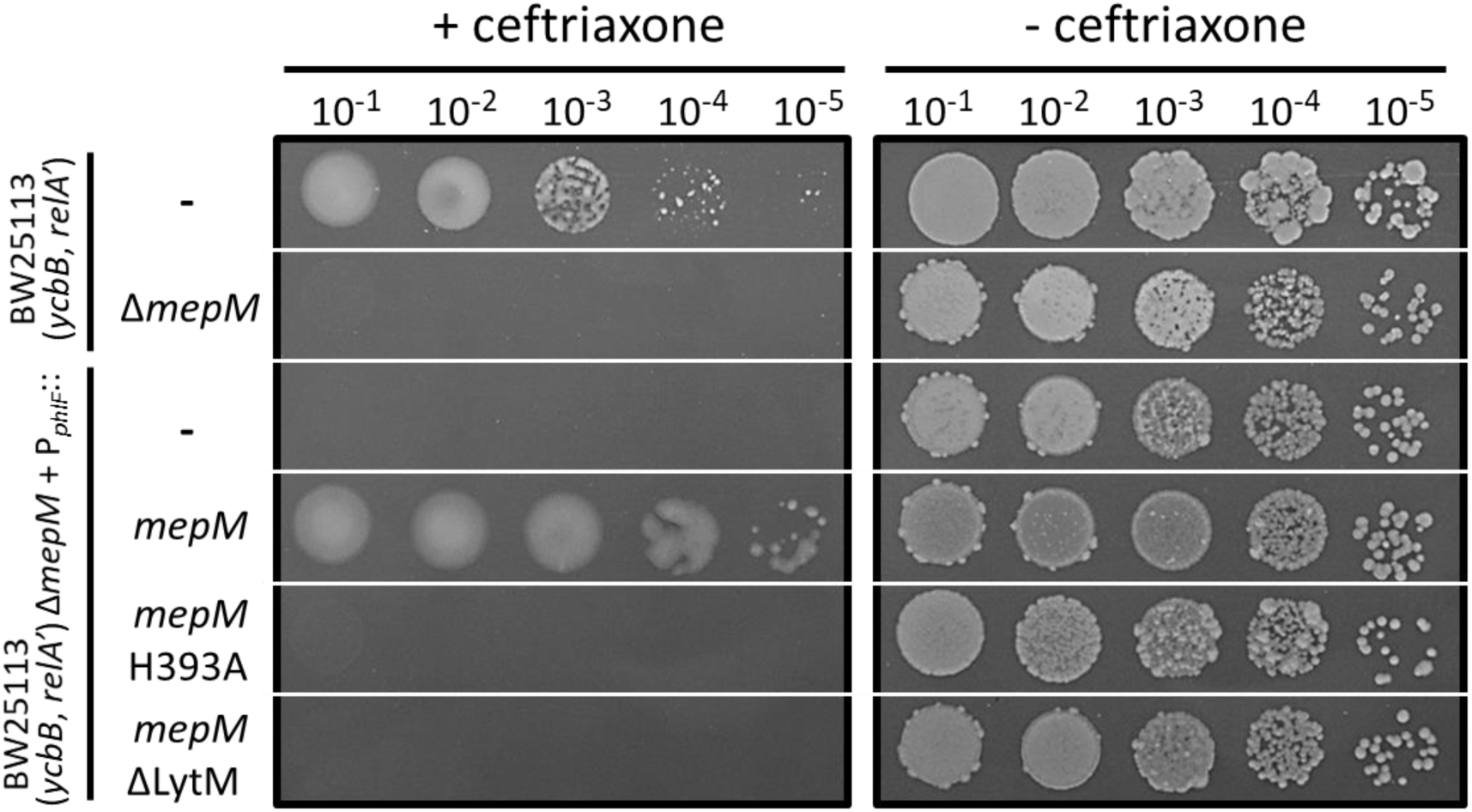
MepM endopeptidase activity is required for YcbB-mediated β-lactam resistance. Growth of BW25113(*ycbB*, *relA’)*, BW25113(*ycbB*, *relA’) ΔmepM*, and its derivatives harboring plasmids encoding MepM, MepM H^393^A, and MepM ΔLytM were tested in the presence of ceftriaxone at 8 μg/ml (+ ceftriaxone) or in the absence of the drug (-ceftriaxone) in BHI agar plates supplemented with 40 μM IPTG and 1% L-arabinose for induction of *ycbB* and *relA’*, respectively. The genes encoding MepM, MepM H^393^A, and MepM ΔLytM were inserted into the vector pHV9 under the control of the *P_phlF_* promoter, which is inducible by 2,4-diacetylphloroglucinol (DAPG). Basal level of expression of *mepM* under the control of *P_phlF_* was sufficient to restore ceftriaxone resistance in the absence of the inducer.

### The essential role of the endopeptidase activity of MepM in β-lactam resistance is not restricted to the transition from 4→3 to 3→3 cross-links triggered by the induction of *ycbB* and *relA’*

The experiment reported above did not rule out the possibility that hydrolysis of 4→3 cross-links by MepM might be transiently essential to enable bypass of PBPs by YcbB, *i.e*. cleavage of 4→3 cross-links by MepM could be initially essential to enable insertion of new PG subunits into the PG network by YcbB. Ultimately, this would lead to replacement of 4→3 by 3→3 cross-links and could then suppress the essential role of 4→3 cross-link cleavage by MepM. According to this hypothesis, MepM would only be essential during the transition between the two modes of PG cross-linking. To test this possibility, we sought a plasmid construct enabling tight regulation of the *mepM* gene. In a first attempt, *mepM* was cloned under the control of the L-rhamnose-inducible promoter *(P_rhaBAD_)* of the pHV30 vector. Complementation of the *mepM* deletion of BW25113(*ycbB*, *relA’)* was obtained both in the presence or absence of L-rhamnose indicating that the un-induced level of *ycbB* afforded by this plasmid construct was too high (Fig. 4A). To address this issue, the level of *mepM* expression was reduced by replacing the sequence containing the translation initiation signal (TIS1) of *mepM* by a weaker translation initiation signal (TIS2). TIS1 (a<u>AAGAGG</u>A<u>GA</u>AAtgacataATG) combined an ATG initiation codon to a “strong” ribosome-binding-site (RBS) with extensive complementarity (underlined) to the 3’ OH extremity of 16S rRNA (5’-A<u>UC</u>A<u>CCU</u>C<u>CUUA</u>-3’OH) (Elowitz and Leibler, 2000). TIS2 (acac<u>AGGA</u>cacttaTTG) combined a TTG initiation codon to an RBS with limited complementarity to 16S rRNA (5’-AUCACC<u>UCCU</u>UA-3’OH) (Hecht et al., 2017; Ringquist et al., 1992; Vellanoweth and Rabinowitz, 1992). In contrast to the results obtained with TIS1, the presence of L-rhamnose was required for β-lactam resistance if *mepM* was expressed under the control of TIS2 (Fig. 4A). L-rhamnose requirement for growth in the presence of ceftriaxone was not abolished by pre-exposure to the inducer (Fig. 4B). These results indicate that the essential role of MepM in β-lactam resistance is not limited to the transition between the two modes of PG cross-linking, *i.e*. from 4→3 to 3→3. Since all D,D-transpeptidases are inhibited by ceftriaxone, these results also indicate that the hydrolytic activity of MepM is essential in conditions in which 4→3 cross-links are not detectable (Hugonnet et al., 2016; Kocaoglu and Carlson, 2015).

**Figure 4.**
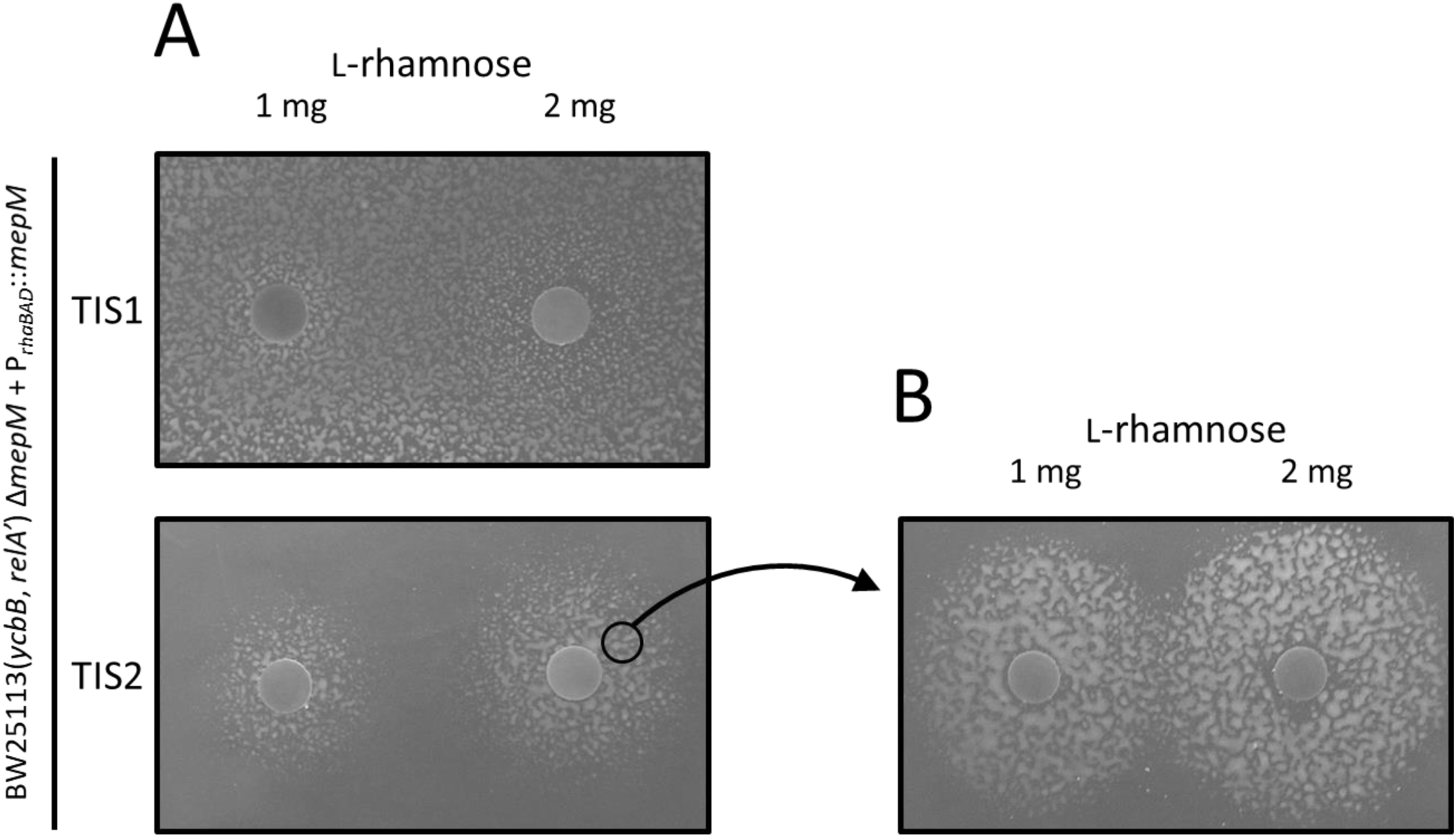
MepM is essential for β-lactam resistance beyond the transition from 4→3 to 3→3 PG cross-linking. The *mepM* gene was expressed under the control of TIS1 or TIS2 (translation initiation) and of *P_rhaBAD_* (the L-rhamnose-inducible promoter of vector pHV30). Growth of BW25113(*ycbB*, *relA’) ΔmepM* harboring the pHV30 derivatives was tested on BHI agar supplemented with ceftriaxone 8 μg/ml, IPTG 40 μM (induction of *ycbB)* and L-arabinose 1% (induction of *relA’)*. (**A**) For TIS2, growth around paper disks containing 1 or 2 mg of L-rhamnose indicated that induction of the expression of *mepM* was required for ceftriaxone resistance. In contrast, a higher level of translation from TIS1 was sufficient for ceftriaxone resistance in the absence of the inducer. (**B**) The experiment was repeated with bacteria pre-exposed to L-rhamnose that were harvested at the vicinity of the disk containing 2 mg of L-rhamnose. Expression of ceftriaxone resistance remained dependent on the presence of L-rhamnose indicating that the requirement for MepM is not transient. The diameter of the growth zones is larger in panel (**B**) than in panel (**A**) as expected from the fact that bacteria in the inoculum used in (**B**) had been grown in the presence of the inducer and already contained MepM. In (**A**) growth is only possible after diffusion of L-rhamnose in the medium prior to the action of ceftriaxone. At a distance from the disk, diffusion was not sufficiently rapid to observe resistance.

### MepM hydrolyzes both 4→3 and 3→3 cross-links *in vitro*

The essential role of the endopeptidase activity of MepM in the context of LDT-mediated PG cross-linking (above) led us to reconsider the specificity of the enzyme. Previous analyses were based on incubation of MepM and lysozyme with an *E. coli* PG preparation containing minute amounts of 3→3 cross-links (Chodisetti and Reddy, 2019; Singh et al., 2012). Analyses of *rp*HPLC profiles revealed that the major dimers containing 4→3 cross-links were digested by MepM but minor peaks corresponding to dimers containing 3→3 cross-links remained unchanged in the presence of the enzyme leading to the conclusion that MepM was specific to 4→3 cross-links (Chodisetti and Reddy, 2019; Singh et al., 2012). To improve the sensitivity of the assay, we purified MepM and reproduced this analysis with a PG preparation of *E. coli* BW25113 grown to stationary phase in minimal medium, which contained a higher proportion of 3→3 cross-links (Fig. 5A, upper panel.) The muropeptides corresponding to the indicated peaks are shown in Fig. 5B (see supplementary Fig. S2 for determination of the structure of muropeptides by mass spectrometry). Full digestion of all dimers was observed upon incubation of this PG preparation with MepM (5 μM) indicating that the endopeptidase hydrolyzes both 4→3 and 3→3 cross-links to completion (Fig. 5A, lower panel). Incubation of the PG preparation with lower concentrations of MepM led to partial hydrolysis of the dimers (Fig. 5C). Comparison of the relative abundance of muropeptides based on the integration of peak areas in the chromatograms showed the expected increase in monomers upon digestion of dimers (Fig. 5D). The concentrations of MepM required for hydrolysis of half of the muropeptides containing 4→3 and 3→3 cross-links were 0.4 μM and 0.7 μM, respectively, revealing similar apparent hydrolysis efficacies for the two types of cross-links under the assay conditions (Fig. 5D). These results indicate that hydrolysis of 3→3 cross-links may account for the essential role of MepM in conditions in which the L,D-transpeptidase activity of YcbB fully replaces the D,D-transpeptidase activity of the PBPs, as inferred from expression of ceftriaxone resistance by BW25113(*ycbB*, *relA’)* but not by its *ΔmepM* derivative (Fig. 2B).

**Figure 5.**
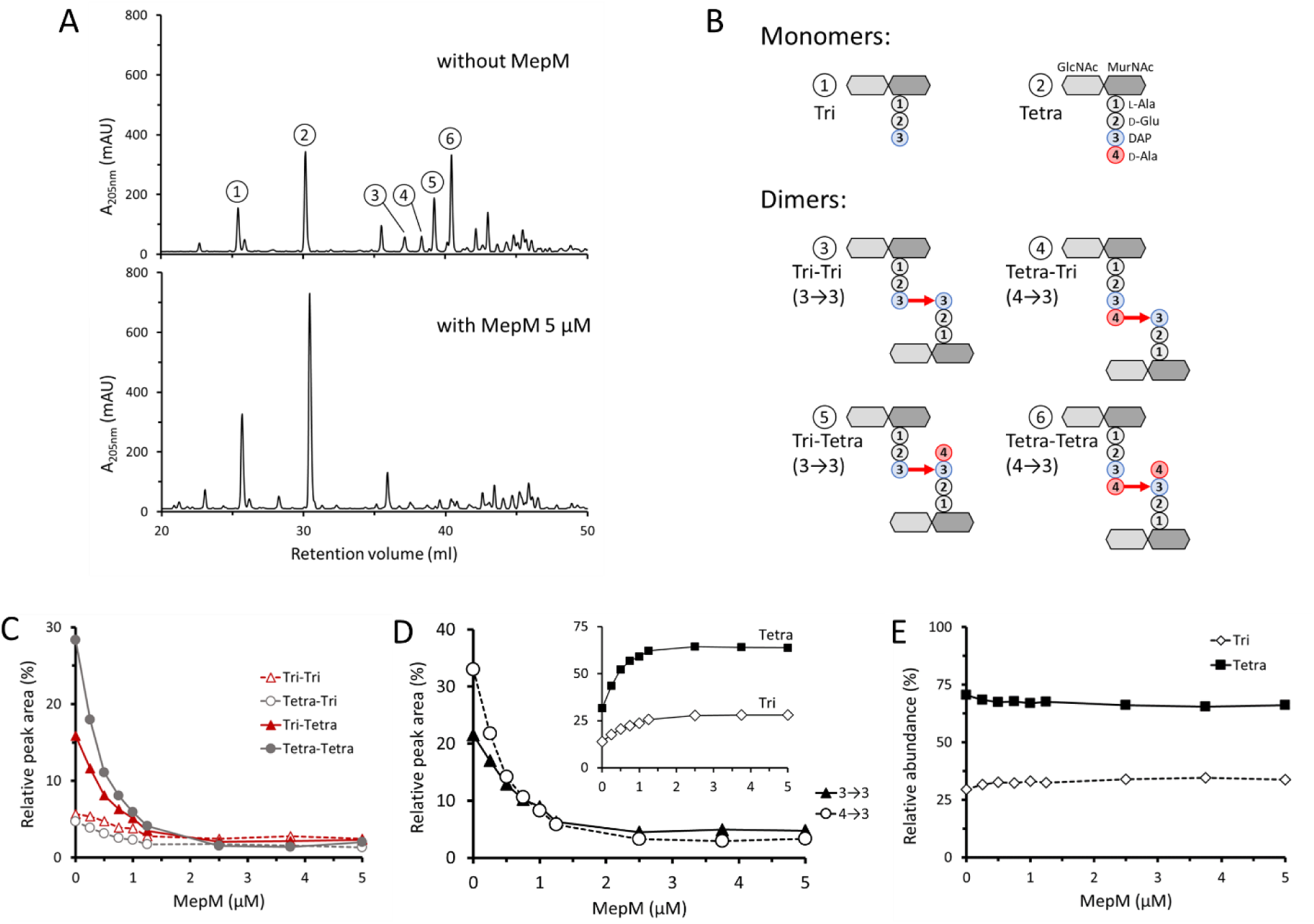
Hydrolysis of 4→3 and 3→3 cross-links by purified MepM. (**A**) *rp*HPLC chromatograms of sacculi isolated from BW25113 grown in minimal medium to stationary phase and digested by lysozyme (upper panel) or by lysozyme and 5 μM MepM (lower panel). Absorbance was monitored at 205 nm (mAU, milliabsorbance unit). (**B**) Structure of the muropeptides as determined by mass spectrometry (supplementary Fig. S2). (**C**) Hydrolysis of the four types of dimers by MepM. Sacculi were incubated with lysozyme and MepM at various concentrations. The relative abundance of the muropeptides was estimated by calculating the relative peak areas. (**D**) Hydrolysis of dimers containing 4→3 and 3→3 cross-links by MepM. The relative peak areas of Tri-Tri and Tri-Tetra containing 3→3 cross-links and that of Tetra-Tri and TetraTetra containing 4→3 cross-links were combined. The inset shows variations in the relative peak areas of the Tri and Tetra monomers. (**E**) Relative abundance of Tri and Tetra stems in all muropeptides.

The sum of the relative proportion of tripeptide and tetrapeptide stems in all monomers and dimers did not vary upon addition of MepM (Fig. 5E). This observation indicates that MepM did not hydrolyze D-Ala^4^ from tetrapeptide monomers or from tetrapeptide stems located in the acceptor position of dimers. Thus, MepM did not display any L,D-carboxypeptidase activity.

**Figure S2.**
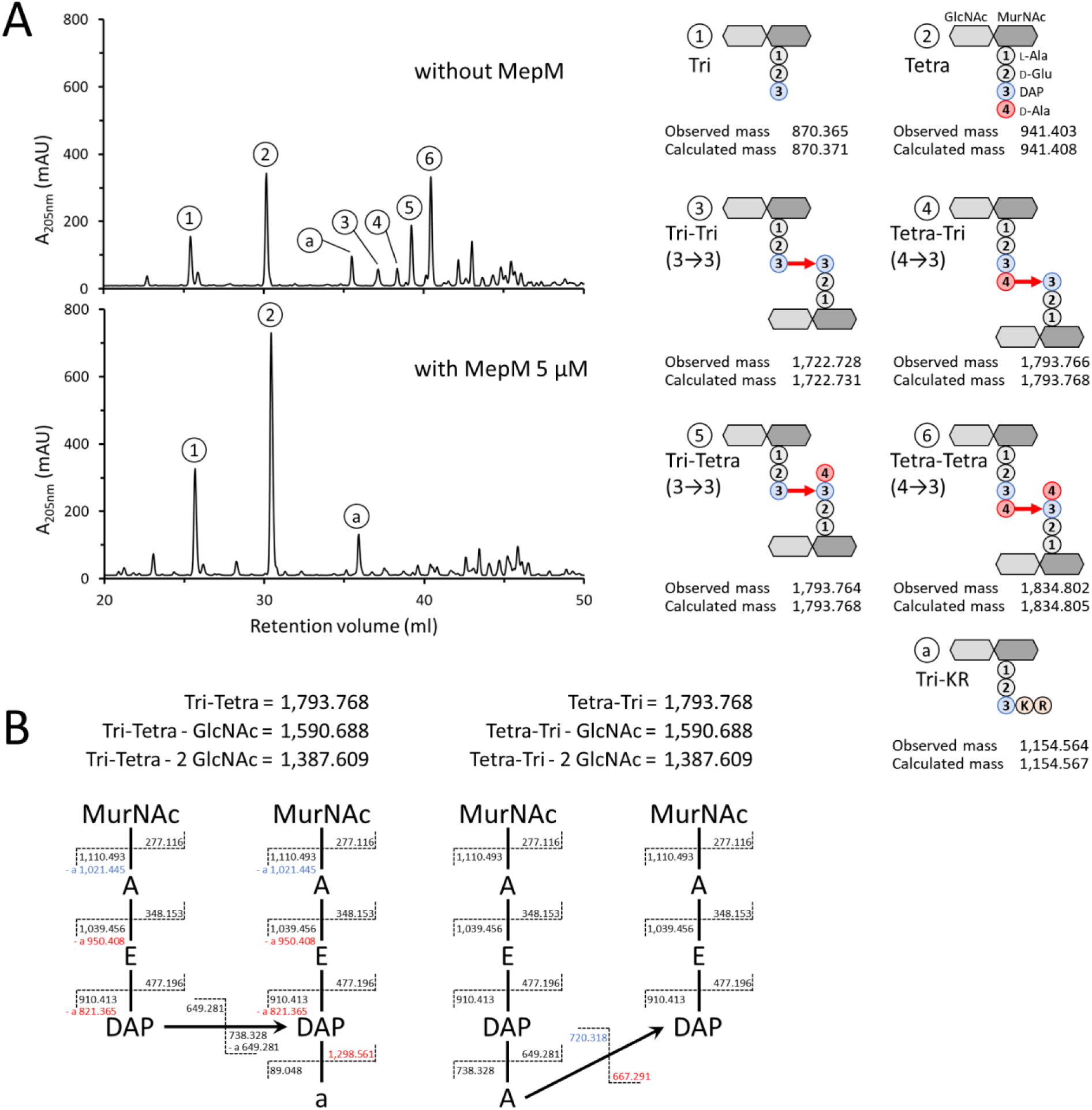
(**A**) Mass spectrometry analysis of muropeptides obtained by digestion of sacculi from BW25113 with lysozyme (upper panel) or lysozyme plus MepM (lower panel). The observed and calculated monoisotopic mass are indicated in Dalton. Peak a corresponds to a disaccharide-tripeptide substituted by a Lys-Arg (KR) dipeptide originating from digestion of the covalently-bond Braun lipoprotein by trypsin (Magnet et al., 2008). (**B**) Discrimination of isomers containing 3→3 (Tri-Tetra) and 4→3 (Tetra-Tri) cross-links by tandem mass spectrometry. All fragments lost both GlcNAc molecules. Fragments that are specific of each isomer are shown in red. Fragments specific of an isomer but which can also be found in the other isomer following loss of a water molecule are shown in blue. Mass of fragments is shown in Dalton. A, L-Ala or D-Ala; a, C-terminal D-Ala; E, D-Glu; DAP, diaminopimelic acid.

### Design of an assay to investigate the redundancy of endopeptidases required for YcbB-mediated β-lactam resistance

Single deletion of endopeptidase genes revealed four phenotypes (above, Fig. 2B). (i) Deletion of *mepK* was not compatible with production of YcbB. (ii) Deletion of *mepM* abolished YcbB-mediated ceftriaxone resistance. (iii) Deletion of *mepS* impaired growth in the presence of ceftriaxone. (iv) Deletion of *mepA*, *mepH*, *dacB*, *pbpG*, or *ampH* had no impact on growth in the presence of ceftriaxone. The absence of any phenotypic alteration associated with the individual deletion of the latter genes does not necessarily imply that the corresponding endopeptidases are unable to participate in the hydrolysis of 3→3 cross-links. Indeed, the function of these enzymes may be redundant. Alternatively, their level of production may be insufficient under the tested growth conditions. To investigate these possibilities, each of the eight endopeptidase genes was independently cloned under the control of the “strong” TIS1 translation initiation signal downstream from the *P_rhaBAD_* promoter of the vector pHV30 in order to modulate the level of endopeptidase production based on induction by L-rhamnose. The plasmids were introduced into BW25113(*ycbB*, *relA’) ΔmepM* and growth of the resulting strains was tested in the presence or absence of L-rhamnose and in the presence or absence of ceftriaxone in all combinations (Fig. 6).

**Figure 6.**
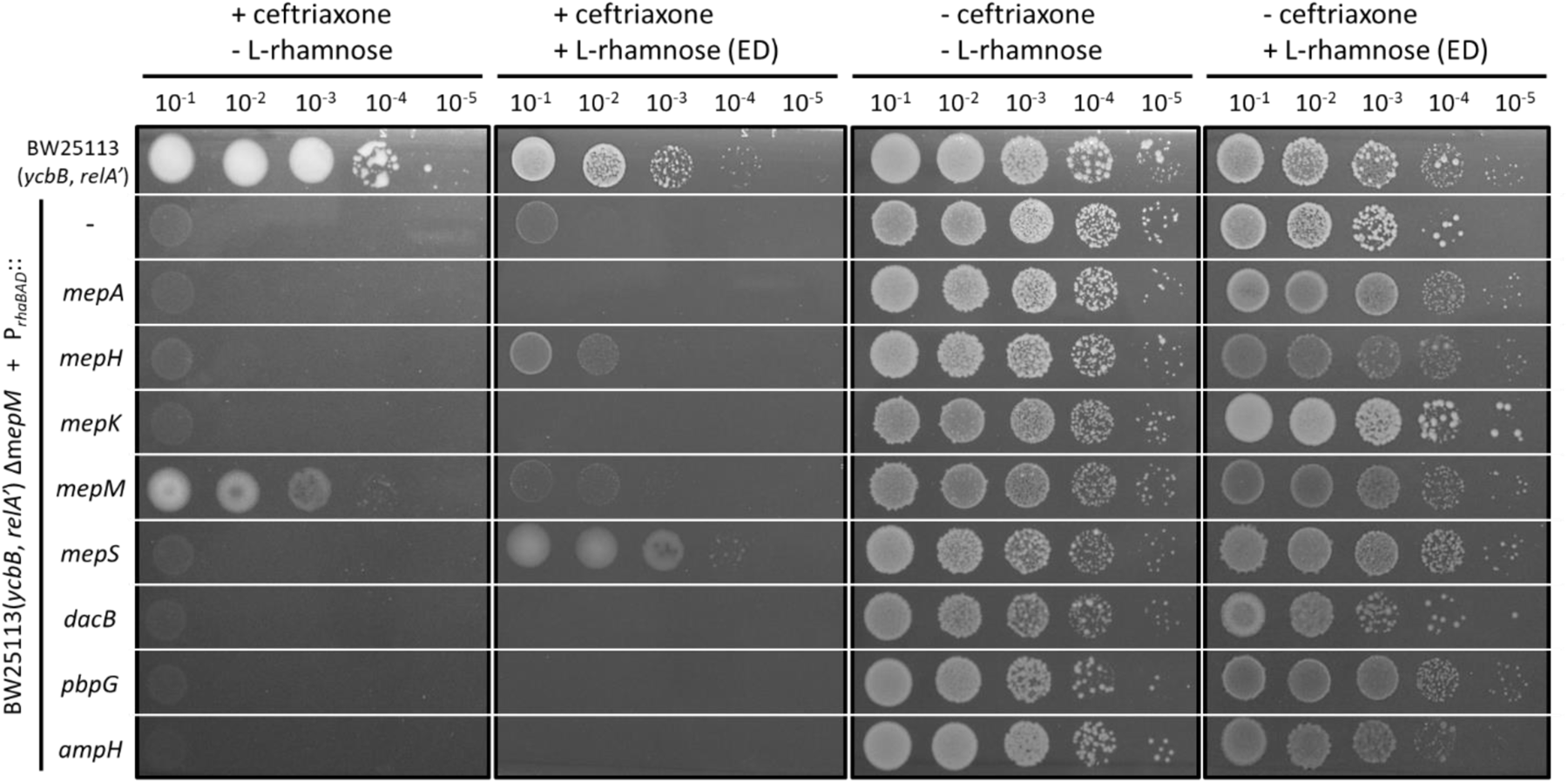
Complementation of the *mepM* deletion by plasmids encoding L-rhamnose-inducible copies of the eight endopeptidase genes. Functional complementation of the *mepM* deletion in BW25113(*ycbB*, *relA’) ΔmepM* was performed with the pHV30 vector or recombinant plasmids encoding each of the eight endopeptidases under the control of the *P_rhaBAD_* promoter. Induction of endopeptidase (ED) genes was performed with 0.2% L-rhamnose in the presence or absence of 8 μg/ml ceftriaxone. BHI agar plates contained 40 μM IPTG and 1% L-arabinose for induction of *ycbB* and *relA’*, respectively.

### Overexpression of *mepM* is toxic in the presence of ceftriaxone

The assay described above revealed that the basal level of expression of the plasmid copy of *mepM* in the absence of the inducer was sufficient to restore growth of BW25113(*ycbB*, *relA’) ΔmepM* in the presence of ceftriaxone (Fig. 6). Induction of the *mepM* gene by L-rhamnose prevented growth in the presence of ceftriaxone but not in the absence of the drug. These results suggest that overproduction of MepM inhibits growth by cleavage of 3→3 cross-links if 4→3 cross-links are absent due to the inactivation of the PBPs by ceftriaxone.

### Overexpression of *mepS* complements the *mepM* deletion for expression of YcbB-mediated β-lactam resistance

MepS restored growth of BW25113(*ycbB*, *relA’) ΔmepM* only in the presence of the inducer (Fig. 6). Thus, MepM and MepS have overlapping functions although overproduction of MepS was required to compensate for the absence of MepM. As mentioned above (Fig. 2B), deletion of *mepS* impaired but did not abolish ceftriaxone resistance in BW25113(*ycbB*, *relA’)*. Together these results indicate that expression of *mepS* in its native chromosomal environment contributes to resistance but the level of its expression is not sufficient to compensate for the absence of MepM.

### Partial complementation of the *mepM deletion* by *mepH*

The plasmid encoding MepH partially restored growth of BW25113(*ycbB*, *relA’) ΔmepM* on ceftriaxone only in the presence of L-rhamnose (Fig. 6). This result indicates that MepH, like MepS, replaces MepM for the expression of ceftriaxone resistance if MepH is overproduced.

### Purified MepS and MepH hydrolyze 4→3 and 3→3 cross-links (endopeptidase activity) and the DAP-D-Ala bond of tetrapeptide stems (L,D-carboxypeptidase activity)

Complementation of *ΔmepM* by overproduction of MepS and MepH prompted us to evaluate the specificity of these enzymes, as described above for MepM. MepH and MepS both hydrolyzed 4→3 and 3→3 cross-links (Fig. 7A and B). MepS showed no preference for 4→3 or 3→3 cross-links while MepH displayed a strong preference for 4→3 cross-links. The weak hydrolytic activity of MepH on 3→3 cross-links may account for the fact that the overproduction of MepH can only partially compensate for the absence of MepM (Fig. 6, above). Both MepS and MepH displayed L,D-carboxypeptidase activity leading to rapid conversion of tetrapeptide stems into tripeptide stems (Fig. 7A and B).

**Figure 7.**
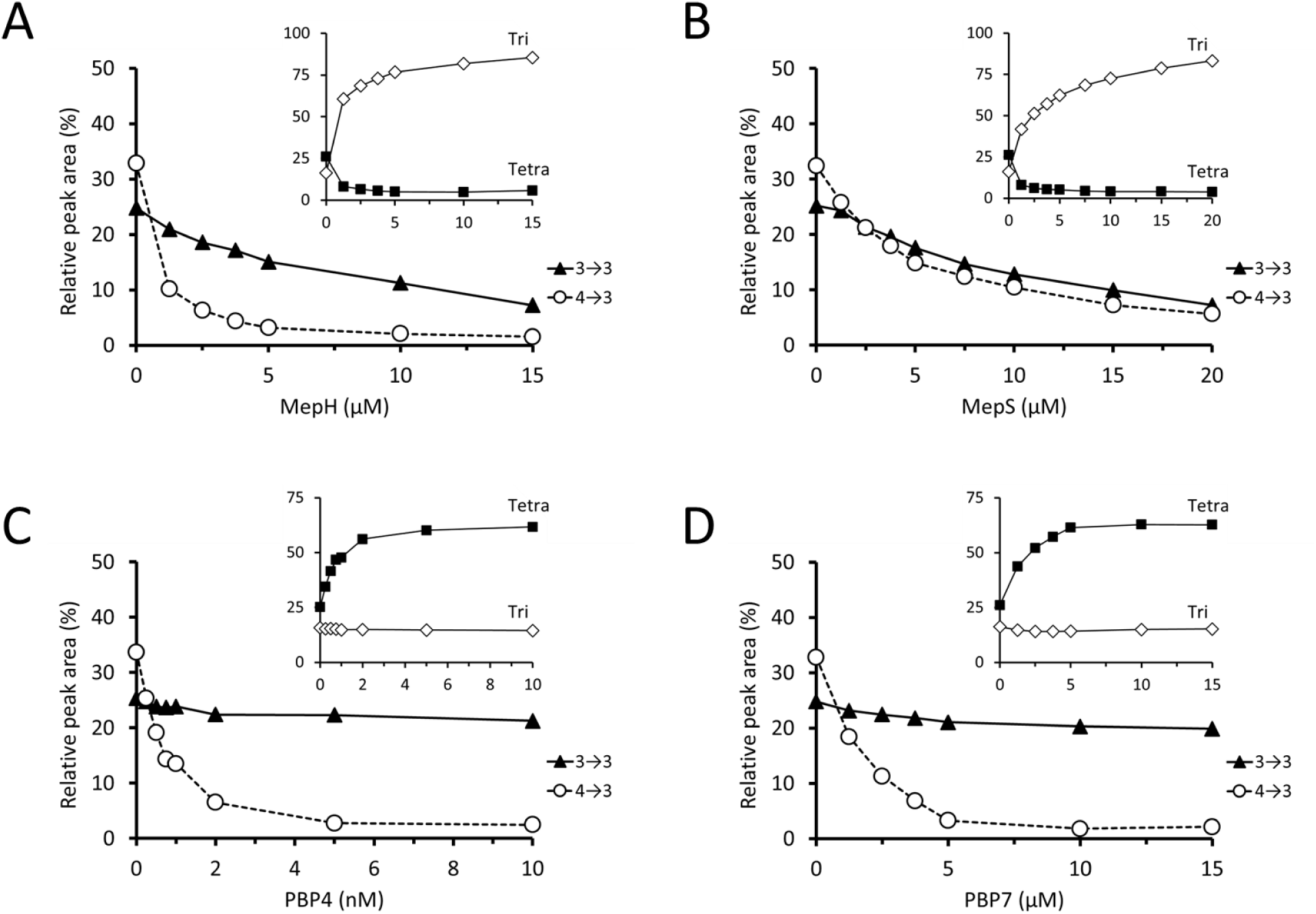
Hydrolysis of 4→3 and 3→3 cross-links by purified endopeptidases. Sacculi were incubated with lysozyme and purified MepH (**A**), MepS (**B**), PBP4 (**C**), and PBP7 (**D**). The relative peak areas of Tri-Tri and Tri-Tetra containing 3→3 cross-links and that of Tetra-Tri and Tetra-Tetra containing 4→3 cross-links were combined. MepH preferentially hydrolyzed dimers containing 4→3 cross-links. MepS hydrolyzed dimers containing 4→3 and 3→3 cross-links with similar efficacies. PBP4 and PBP7 only hydrolyzed dimers containing 4→3 cross-links. The insets show variations in the relative peak areas of the Tri and Tetra monomers. MepH and MepS displayed L,D-carboxypeptidase activity, but not PBP4 and PBP7.

### MepA and MepK do not compensate the absence of MepM

In spite of the fact that MepA and MepK were previously shown to cleave 3→3 cross-links (Chodisetti and Reddy, 2019; Engel et al., 1992), growth of BW25113(*ycbB*, *relA’) ΔmepM* on ceftriaxone was not restored by overproduction of these enzymes (Fig. 6). Thus, there was not a strict correlation between the ability of the endopeptidases to hydrolyze 3→3 cross-links *in vitro* and their ability to restore growth of BW25113(*ycbB*, *relA’) ΔmepM*. This absence of correlation was particularly striking for MepK since this endopeptidase, which preferentially hydrolyze 3→3 cross-links *in vitro* (Chodisetti and Reddy, 2019), did not complement the *mepM* deletion although it was required for growth of BW25113(*ycbB*, *relA’)* expressing the *ycbB* L,D-transpeptidase gene.

These results indicate that functional properties of the endopeptidases, beyond their mere hydrolytic specificity, are relevant to the bypass of the D,D-transpeptidase activity of PBPs by the L,D-transpeptidase activity of YcbB. These properties may include the interaction of the endopeptidases with other proteins that regulate their spatiotemporal activity (see discussion section).

### Endopeptidases of the PBP family do not compensate the absence of MepM

Complementation of the *mepM* deletion was not observed for PBP4, PBP7, and AmpH both in the presence or absence of induction of the corresponding genes by L-rhamnose (Fig. 6). There is a caveat for these endopeptidases since they are potentially inhibited by ceftriaxone. To address this issue, the complementation test was repeated with ampicillin, cefsulodin, and aztreonam, which were reported to exhibit different selectivities for inhibition of the PBPs (Henderson et al., 1997; Kocaoglu and Carlson, 2015). Plasmids encoding PBP4, PBP7, and AmpH did not restore growth of BW25113(*ycbB*, *relA’) ΔmepM* in the presence of ampicillin, cefsulodin, and aztreonam (supplementary Fig. S3) confirming that these endopeptidases are unable to compensate for the absence of MepM. PBP4 and PBP7 were purified and shown to only cleave 4→3 cross-links (Fig. 7C and D). Thus, the absence of complementation of the *mepM* deletion by the plasmids encoding these PBPs can be accounted for by their lack of hydrolytic activities on 3→3 cross-linked dimers. PBP4 and PBP7 did not display L,D-carboxypeptidase activity.

**Figure S3.**
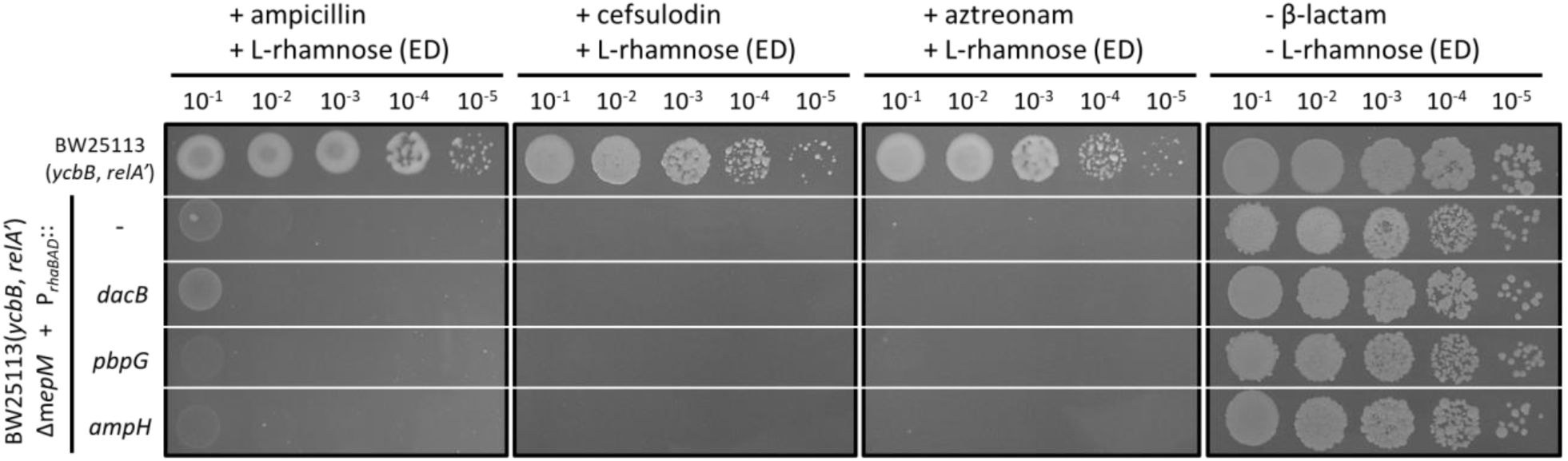
Complementation of the *mepM* deletion by endopeptidases of the PBP family. Functional complementation of the *mepM* deletion of BW25113(*ycbB*, *relA’) ΔmepM* was performed with the pHV30 vector or recombinant plasmids encoding PBP4, PBP7, and AmpH under the control of the *P_rhaBAD_* promoter. Induction of endopeptidase (ED) genes was performed with 0.2% L-rhamnose in the presence or absence of 16 μg/ml ampicillin, 32 μg/ml cefsulodin, or 8 μg/ml aztreonam. BHI agar plates contained 40 μM IPTG and 1% L-arabinose for induction *ycbB* and *relA’*, respectively.

### Minimal complement of endopeptidases required for growth in the context of the formation of 4→3 cross-links by PBPs

Previous analyses based on multiple deletions showed that genes encoding endopeptidases belonging to the PBP family (PBP4, PBP7, and AmpH) are collectively dispensable (Denome et al., 1999). Independently, deletion of the genes encoding MepH, MepM, and MepS in various combinations revealed that at least one of these endopeptidases was essential (Singh et al., 2012). Here, we extend these analyses to the full complement of the eight endopeptidase genes.

Serial deletions of endopeptidase genes were introduced into the chromosome of *E. coli* BW25113 *ΔrelA* generating the lineages depicted in supplementary Fig. S4. This approach culminated in the construction of a viable derivative of BW25113 *ΔrelA*, designated Δ7EDs (lineage 5 in supplementary Fig. S4), which retained only one of the eight endopeptidase genes (*mepM*). Thus, MepM alone was necessary and sufficient to support bacterial growth in the context of the 4→3 mode of cross-linking.

Our next objective was to determine whether deletion of *mepM* could be complemented by overproduction of other endopeptidases. To address this question, the *mepM* gene was cloned under the L-rhamnose-inducible promoter of vector pHV30 and introduced into the Δ7EDs strain. The chromosomal copy of *mepM* was deleted from the resulting strain leading to strain Δ8EDs pHV53(*mepM*), which was dependent upon the presence of L-rhamnose for growth (Fig. 8A). The plasmids enabling L-arabinose-inducible expression of the eight endopeptidase genes (above) were introduced in the Δ8EDs pHV53(*mepM*) strain to determine which endopeptidase could functionally replace MepM (Fig. 8B). Induction by L-arabinose of the genes encoding MepM, MepH, MepS, and PBP7 suppressed the requirement for L-rhamnose for growth. These results indicate that a single endopeptidase, MepM, MepH, MepS, or PBP7, is potentially sufficient for growth in the context of the 4→3 mode of cross-linking. Except for MepM, this required overproduction of the enzymes following induction of the *P_araBAD_* promoter of the recombinant plasmids.

**Figure 8.**
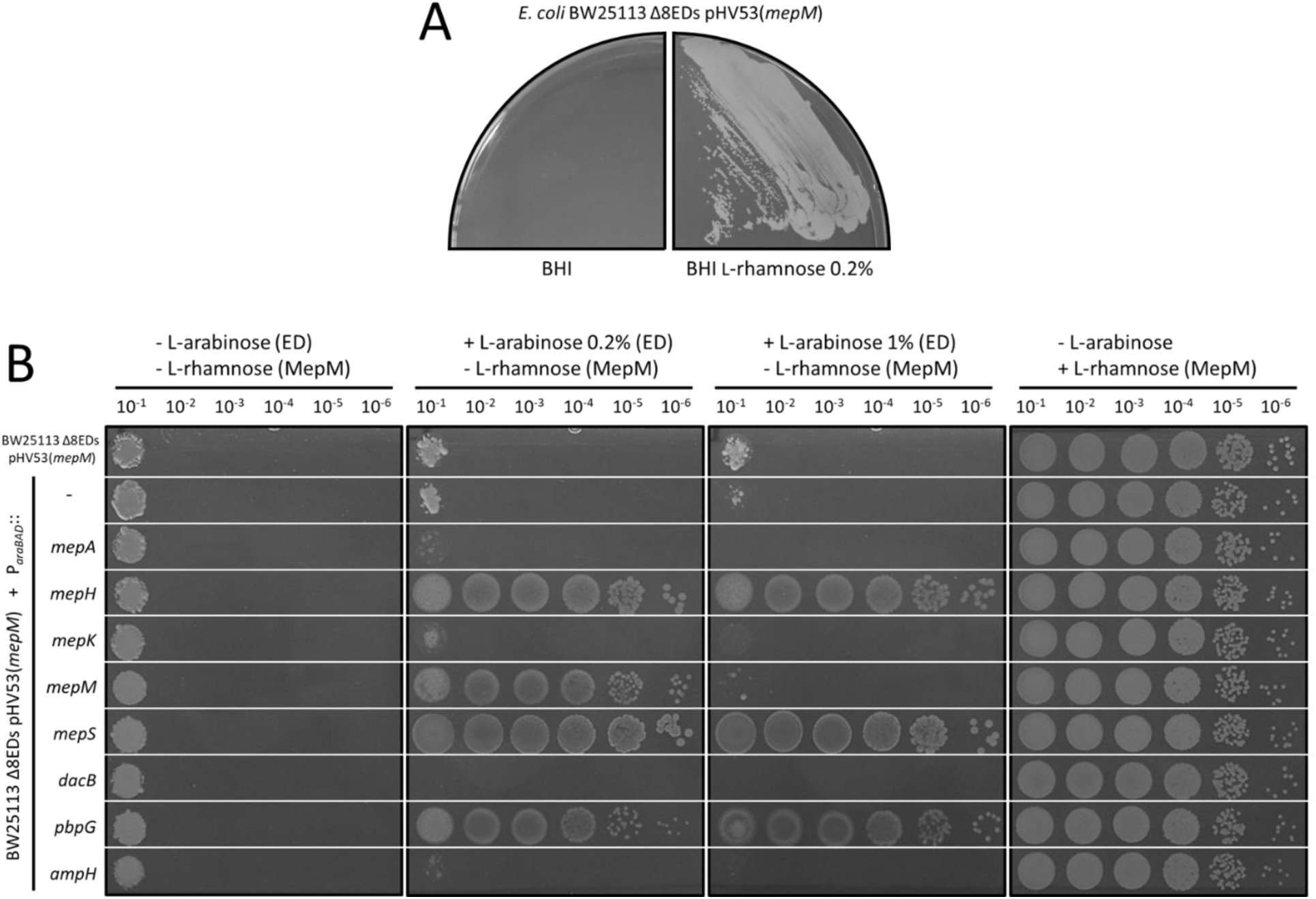
Minimal complement of endopeptidases required for growth in the context of the formation of 4→3 cross-links by PBPs. (**A**) BW25113 Δ8EDs harboring a plasmid carrying the *mepM* gene under the control of the *P_rhaBAD_* L-rhamnose-inducible promoter with the “weak” TIS2 translation initiation signal (plasmid pHV53) was grown on BHI agar in the absence or presence of 0.2% L-rhamnose. Growth was dependent upon induction of the *mepM* copy carried by pHV53. (**B**) The plating efficiency assay was performed with derivatives of BW25113 Δ8EDs pHV53(*mepM*) harboring the vector pHV7 or recombinant plasmids carrying each of the eight endopeptidase genes under the control of the *P_araBAD_* promoter. In this assay, functional replacement of MepM is detected based on growth in media containing 0.2% or 1% L-arabinose for expression of the endopeptidase gene carried by vector pHV7, while by-passing the requirement for induction of the *mepM* copy of pHV53 by L-rhamnose. Complementation was observed with both concentrations of inducer for *mepH, mepS*, and *pbpG*. Overproduction of *mepM* encoded by the pHV7 derivative in the presence of the high dose of L-arabinose (1%) was lethal. The right panel presents the growth control performed in the presence of 0.2% L-rhamnose for induction of the *mepM* copy carried by pHV53.

**Figure S4.**
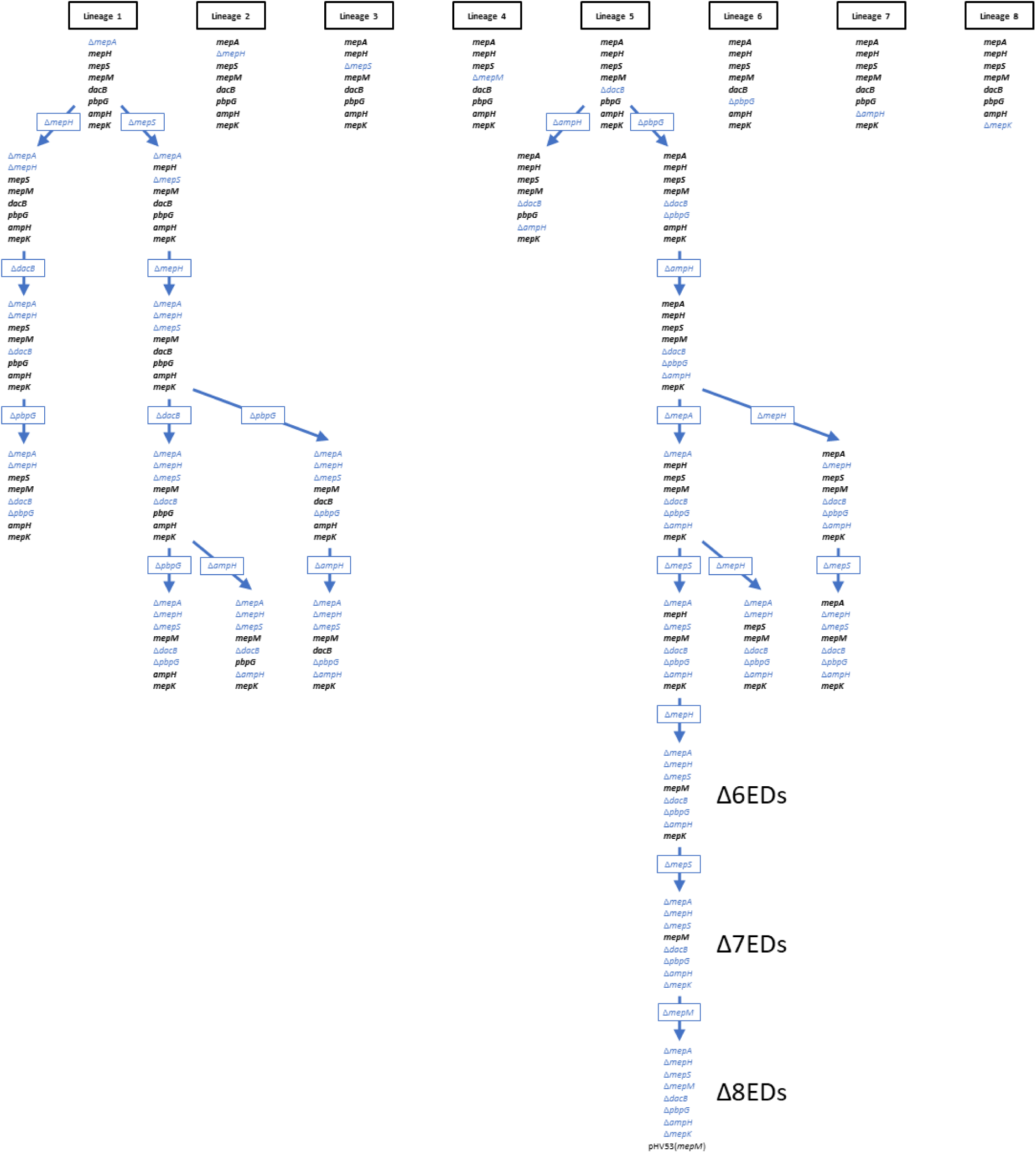
Parallel and serial deletion of endopeptidase genes in *E. coli* BW25113 *ΔrelA*. Deletions indicated in blue were introduced by the procedure of Datsenko and Wanner (Datsenko and Wanner, 2000). The remaining endopeptidase genes are indicated in black. The presence of all deletions was verified by PCR at each steps. The genomes of the strains retaining *mepM* and *mepK* (Δ6EDs) or only *mepM* (Δ7EDs) were re-sequenced and no compensatory mutation was detected.

### MepA and MepS compensate for the absence of MepK when *ycbB* is induced

The basal production of the YcbB L,D-transpeptidase encoded by plasmid pKT2(*ycbB*) in the absence of induction was found to be lethal in a derivative of BW25113 lacking *mepK* (above). To investigate the possibility that MepK might be replaced by another endopeptidase, the *ycbB* gene was cloned under the control of the *P_araBAD_* promoter of vector pHV7 to obtain a lower level of expression of the L,D-transpeptidase gene. The resulting plasmid, pHV63(*ycbB*) was successfully introduced into the BW25113 *ΔmepK* strain indicating that the basal level of expression of *ycbB* in the absence of induction was compatible with the absence of *mepK*. The disk diffusion assay revealed a clear zone around the disk containing L-arabinose indicating that induction of *ycbB* in the *ΔmepK* background prevented bacterial growth (Fig. 9A). Plasmids for expression of each of the eight endopeptidases under the control of the *P_rhaBAD_* promoter (above) were introduced in this strain (Fig. 9B). Bacterial growth was observed in conditions of induction of *ycbB* by L-arabinose and of genes encoding MepA and MepS by L-rhamnose. This result indicates that the essential role of MepK for PG polymerization mediated by the YcbB L,D-transpeptidase was bypassed by overproduction of MepA or MepS.

**Figure 9.**
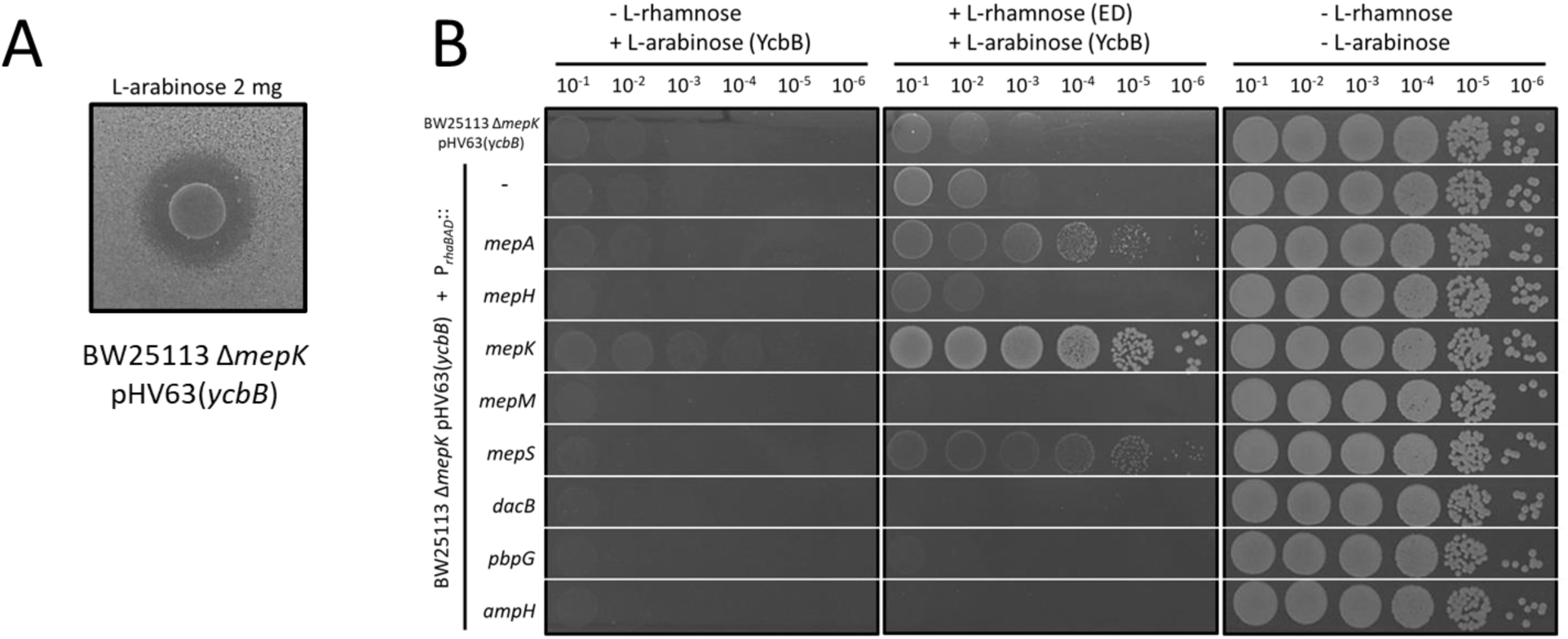
Complementation of the *mepK* deletion by plasmids encoding L-rhamnose-inducible copies of the eight endopeptidase genes. (**A**) Induction of *ycbB* under the control of the *P_araBAD_* promoter in BW25113 *ΔmepK* pHV63(*ycbB*) was studied by the disk diffusion assay. The clear zone around the disk containing L-arabinose indicates that production of YcbB inhibited growth. (**B**) Functional complementation of the *mepK* deletion of BW25113 *ΔmepK* pHV63(*ycbB*) was performed with the pHV30 vector or recombinant plasmids encoding each of the eight endopeptidases under the control of the *P_rhaBAD_* promoter. Induction of *ycbB* and of endopeptidase (ED) genes was performed with 0.2% L-arabinose and 1% L-rhamnose, respectively. BHI agar plates contained chloramphenicol (20 μg/ml) to counter-select loss of pHV63(*ycbB*).

### Complementation of the *mepK* deletion by catalytically inactivated endopeptidases

Since overproduction of MepK, MepS, and MepA were found to complement the chromosomal deletion of the *mepK* gene (above, Fig. 9) we focused on these three endopeptidases. Plasmids encoding catalytically inactive MepK H^133^A, MepS C^94^A, and MepA H^113^A were used to determine whether the endopeptidase activity of MepK, MepA, and MepS was required to compensate for the chromosomal deletion of *mepK*. Overproduction of the endopeptidases was tested in the *ΔmepK* background (single-deletion mutant retaining all chromosomal endopeptidase genes except *mepK)* and in the Δ7EDs background (seven-deletion mutant retaining only *mepM)* (Fig. 10). Overproduction of MepK but not MepK H^133^A was essential for growth in both backgrounds indicating that the catalytic activity of the endopeptidase was essential. Overproduction of MepS restored growth in both backgrounds but complementation by MepS C^94^A was only observed in the *ΔmepK* single-deletion background. Since the periplasmic protease Prc hydrolyzes MepS (Singh et al., 2015) overproduction of MepS C^94^A may saturate the protease enabling sufficient chromosomally-encoded MepS to escape hydrolysis and support growth. Likewise, saturation of the Prc protease is likely to be responsible for the apparent complementation mediated by overproduction of MepA and MepA H^113^A since overproduction of these enzymes restored growth in the ΔmepK single-deletion background but not in the Δ7EDs background. Together these results indicate that MepS is the only endopeptidase that can compensate for the absence of MepK. This required overproduction of MepS. Alternatively, saturation of the Prc protease by overproduction of MepA, MepA H^113^A, or MepS C^94^A prevented hydrolysis of MepS produced at a lower level from the native chromosomal *mepS* gene.

**Figure 10.**
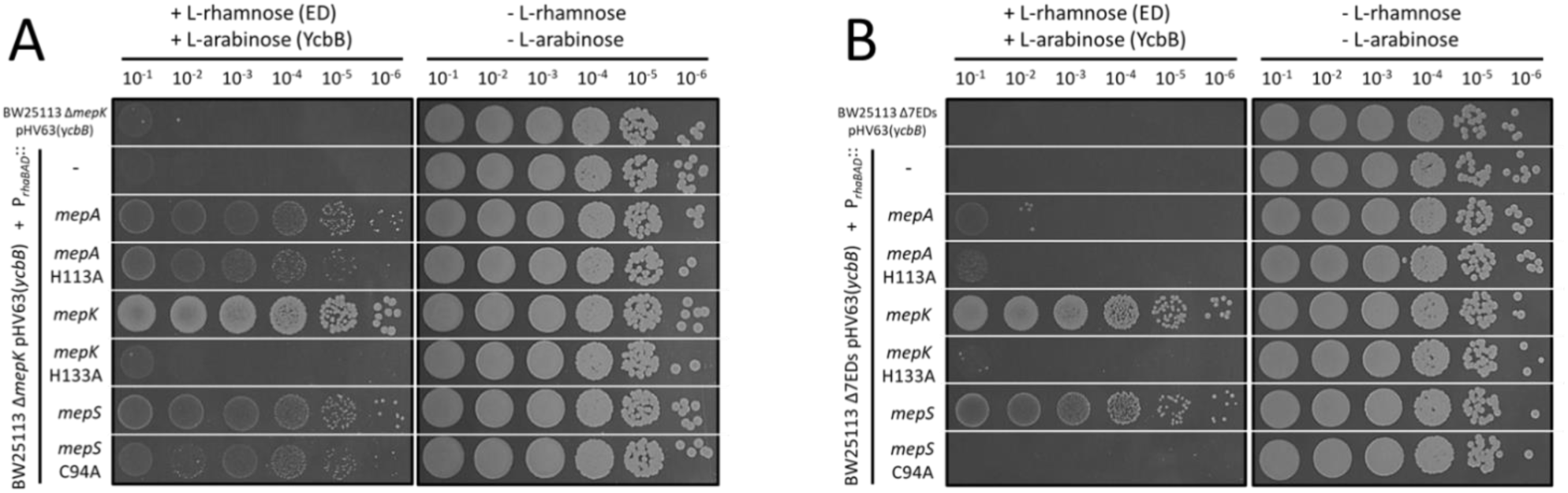
Complementation of *mepK* deletion with catalytically inactivated endopeptidases. (**A**) Functional complementation of the *mepK* deletion of BW25113 *ΔmepK* pHV63(*ycbB*) was performed with the pHV30 vector or recombinant plasmids encoding *mepA, mepK, mepS* or derivatives encoding catalytically inactive endopeptidases under the control of the *P_rhaBAD_* promoter. **(B**) The complementation assay was repeated for BW25113 Δ7EDs pHV63(*ycbB*), which was obtained by deletion of all chromosomal endopeptidase genes except *mepM*. Induction of *ycbB* and of endopeptidase (ED) genes was performed with 0.2% L-arabinose and 1% L-rhamnose, respectively. BHI agar plates contained chloramphenicol (20 μg/ml) to counte-select loss of pHV63(*ycbB*).

### Minimal complement of endopeptidases required for growth in the presence of β-lactams in the context of the exclusive formation of 3→3 cross-links by the YcbB L,D-transpeptidase

As previously described (Hugonnet et al., 2016), induction of *relA’* led to mecillinam resistance in BW25113(*ycbB*, *relA’)* whereas induction of both *ycbB* and *relA’* was required for ampicillin and ceftriaxone resistance (Table 1 and supplementary Fig. S5). Strain BW25113(*ycbB*, *relA’)* Δ6EDs was also resistant to the three β-lactams upon induction of *ycbB* and *relA’* indicating that 6 of the 8 endopeptidase genes were dispensable for expression of β-lactam resistance. In contrast to BW25113(*ycbB*, *relA’)*, the Δ6EDs derivative was resistant to ampicillin and ceftriaxone in the absence of induction of *ycbB* by IPTG. The basal level of *ycbB* expression in the absence of induction was required for resistance since susceptibility to ampicillin and ceftriaxone was observed in the absence of pKT2(*ycbB*). These observations indicate that deletion of 6 of the 8 endopeptidase genes was associated with a decrease in the level of expression of *ycbB* required for β-lactam resistance. In combination with the analysis based on single-gene deletions (Fig. 2B), these results show that MepM and MepK are necessary and sufficient for bacterial growth in conditions in which YcbB is the only functional transpeptidase.

**Table 1.** YcbB-mediated β-lactam resistance in BW25113 derivatives harboring all endopeptidase genes or only *mepM* and *mepK* (Δ6ED)

| Host | Plasmid | Inducer<sup>b</sup> | Inhibition zones (mm)<sup>a</sup> |  |  |
| --- | --- | --- | --- | --- | --- |
|  |  |  | Mec | Amp | Cro |
| BW25113 $\Delta relA$ | | | | | |
|  | pKT2(*ycbB*) pKT8(*relA'*) | None | 24 | 22 | 38 |
|  |  | IPTG (*ycbB*) | 28 | 24 | 39 |
|  |  | Ara (*relA'*) | < 8 | 24 | 39 |
|  |  | IPTG + Ara | < 8 | < 8 | 18 |
| BW25113 $\Delta relA$ $\Delta 6EDs$ | | | | | |
|  | pKT2(*ycbB*) pKT8(*relA'*) | None | 17 | 27 | 41 |
|  |  | IPTG (*ycbB*) | 17 | 28 | 43 |
|  |  | Ara (*relA'*) | < 8 | < 8 | 22 |
|  |  | IPTG + Ara | < 8 | < 8 | 19 |
| BW25113 $\Delta relA$ $\Delta 6EDs$ | | | | | |
|  | pKT8(*relA'*) | None | 17 | 27 | 41 |
|  |  | Ara (*relA'*) | < 8 | 27 | 41 |

<sup>a</sup> The diameter of inhibition zones was determined by the disk diffusion assay around disks containing 10 μg mecillinam (Mec), 10 μg ampicillin (Amp), or 30 μg ceftriaxone (Cro). Examples of the original results are presented in supplementary Fig. S5.
<sup>b</sup> The *ycbB* and *relA'* genes carried by plasmid pKT2 and pKT8 were induced with 40 μM IPTG and 1% L-arabinose (Ara), respectively.

**Figure S5.**
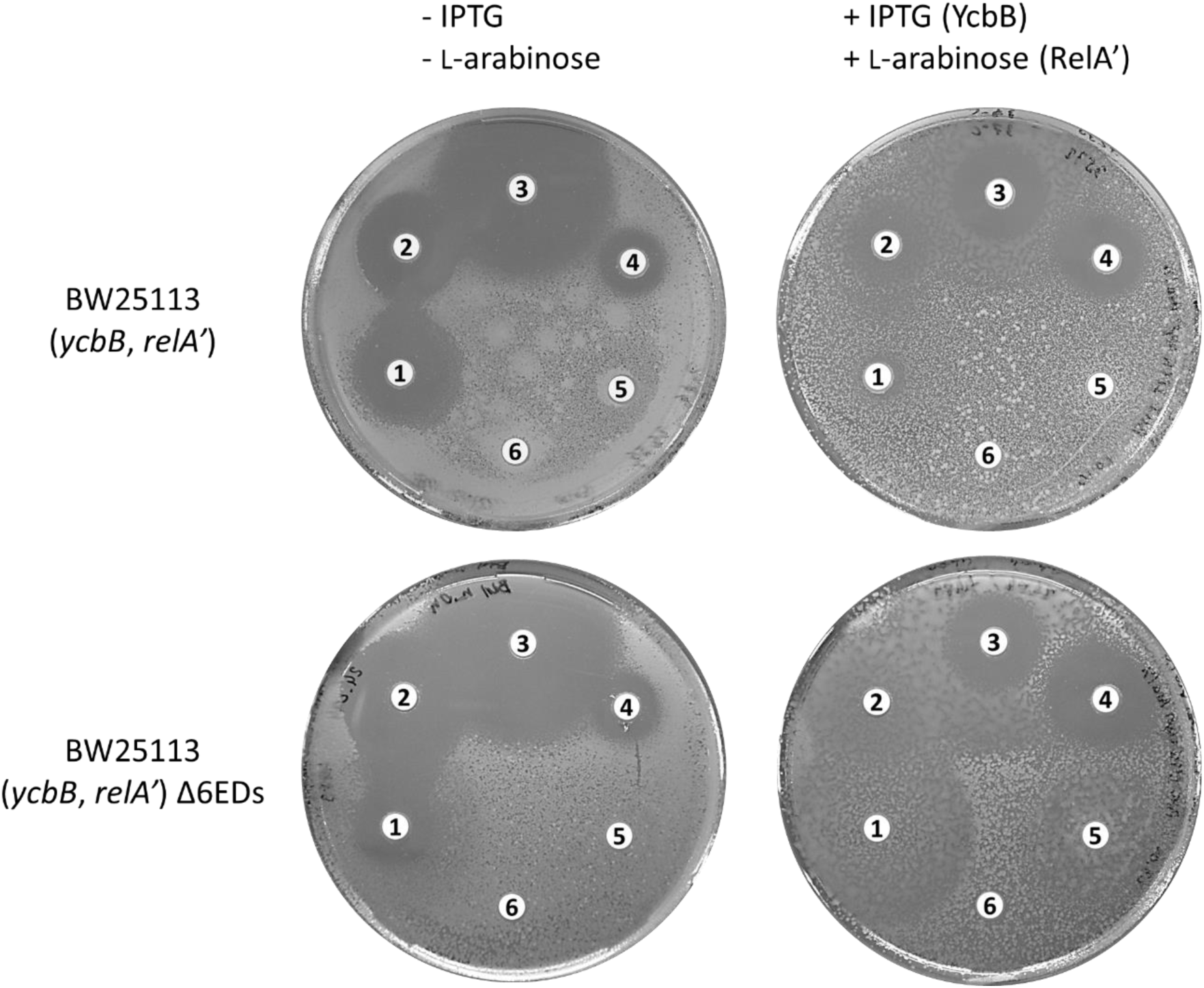
Antibiotic susceptibility testing by the disk diffusion assay. Disks contained 10 μg mecillinam (1), 10 μg ampicillin (2), 30 μg ceftriaxone (3), 30 μg tetracycline (4), 30 μg chloramphenicol (5), and 30 μg kanamycin (6). Plasmids pKT2(*ycbB*) and pKT8(*relA’*) confer resistance to tetracycline and chloramphenicol, respectively. Kanamycin resistance is mediated by the Km^R^ cassette inserted in place of *relA*. Induction of *ycbB* and *relA’* was performed with 40 μM IPTG and 1% L-arabinose.

### Mutations impairing the lytic transglycosylase activity of Slt70 favors YcbB-mediated PG synthesis

Although BW25113(*ycbB*, *relA’)* displays high β-lactam resistance on BHI agar supplemented with IPTG and L-arabinose, the strain was found to remain susceptible to β-lactams in BHI broth supplemented with the same inducers. Mutations leading to expression of β-lactam resistance in liquid medium were sought by selecting mutants derived from *E. coli* BW25113 M1 (Hugonnet et al., 2016). The latter strain overexpresses *ycbB* carried by plasmid pJEH12(*ycbB*) in response to induction by IPTG and overproduces the (p)ppGpp alarmone due to impaired expression of the isoleucine tRNA synthetase gene *ileS* (Hugonnet et al., 2016). Derivatives of BW25113 M1 were selected in BHI broth containing 16 μg/ml ampicillin and 50 μM IPTG. Four independent mutants derived from BW25113 M1 (M1.1 to M1.4) were isolated and shown to grow in liquid medium supplemented with ampicillin and IPTG (Fig. 11A). Whole genome sequencing revealed single mutations all located in the *sltY* gene encoding the Slt70 lytic transglycosylase (Table 2). One of the mutants (M1.3) most probably harbored a null allele of *sltY* since a 7-bp deletion introduced a frame-shift at the 9^th^ codon of the gene. To confirm this conclusion, the *sltY* gene was deleted from the chromosome of BW25113(*ycbB*, *relA’)* strain. The resulting strain, BW25113(*ycbB*, *relA’) ΔsltY*, was also resistant to ampicillin in liquid medium. Growth of BW25113(*ycbB*, *relA’) ΔsltY* in the presence of ampicillin in liquid medium required the presence of IPTG and L-arabinose indicating that overproduction of both the YcbB L,D-transpeptidase and of RelA’ remained essential for β-lactam resistance. Comparison of the resistance phenotype of BW25113(*ycbB*, *relA’*) and its Δ*sltY* derivative on BHI agar revealed that overproduction of YcbB upon induction by IPTG was dispensable for β-lactam resistance in the absence of Slt70 (Fig 11B). However, expression of β-lactam resistance on BHI agar remained dependent upon induction of RelA’ by L-arabinose and upon the presence of pKT2(*ycbB*). These results indicate that loss of Slt70 was essential for expression of YcbB-mediated β-lactam resistance in liquid medium and reduced the level of production of the YcbB L,D-transpeptidase required for expression of resistance on solid medium. This observation suggests that accumulation of uncross-linked glycan chains in the absence of Slt70 may improve the capacity of YcbB to catalyze PG cross-linking accounting for the lower level of expression of *ycbB* required for resistance.

**Figure 11.**
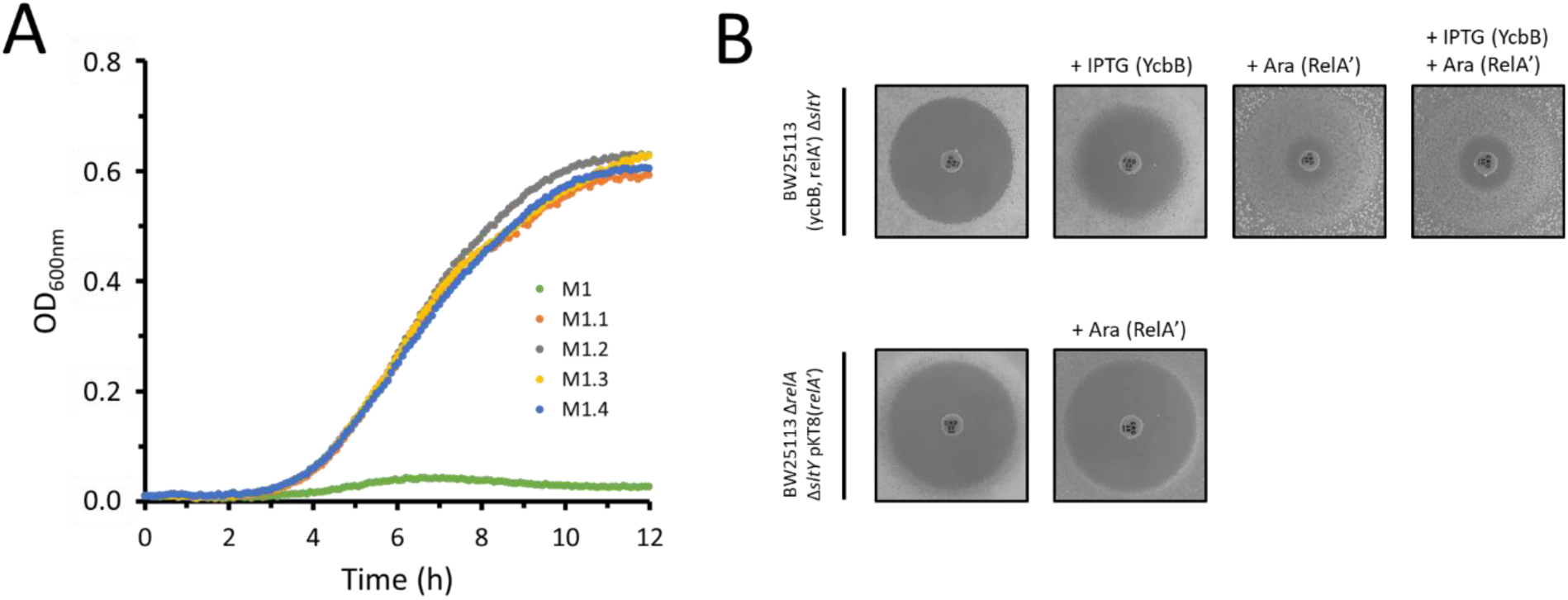
Growth phenotype of *sltY* mutants. (**A**) Growth curves of parental strain M1 and mutants M1.1 to M1.4 selected for expression of ampicillin resistance in BHI broth. The growth medium contained 50 μM IPTG and 16 μg/ml ampicillin. (**B**) Ceftriaxone-resistance depends upon *ycbB* and *relA’* expression. The disk diffusion assay was performed with BW25113 *ΔrelA ΔsltY* harboring pKT2(*ycbB*) and pKT8(*relA’*) or pKT8(*relA’*) only. Induction was performed with 40 μM IPTG and 1% L-arabinose for *ycbB* and *relA’*, respectively. Disks were loaded with 30 μg of ceftriaxone.

**Table 2.** Mutations detected in the *sltY* gene of mutants BW25113 M1.1 to M1.4.

| Mutant | Position | Mutation | Impact<sup>a</sup> |
| --- | --- | --- | --- |
| M1.1 | 4,621,672 | C→T | Gln<sup>375</sup>stop |
| M1.2 | 4,622,297 | Duplication of AGG<br>CAGGG→CAGGAGGG | Insertion of Gly position 584<br>(codon GGN) |
| M1.3 | 4,620,572 | 7-bp deletion<br>CCTGGCGGC→CC | Trp<sup>9</sup>frameshift |
| M1.4 | 4,621,507 | C→T | Arg<sup>320</sup>stop |

<sup>a</sup> Slt70 comprises 645 amino acid residues.

## DISCUSSION

### Endopeptidases are essential for expansion of sacculi

PG polymerization requires a combination of synthetases, the transpeptidases and the glycosyltransferases, in addition to hydrolases that fulfill two essential roles. Since PG is a net-like macromolecule completely surrounding the bacterial cell it is beyond any required experimental demonstration that insertion of new disaccharide-peptide subunits into the growing cell wall requires cleavage of covalent bonds (Höltje and Heidrich, 2001; Vollmer, 2012). The stress-bearing PG being present during the entire cell cycle, it is also obvious that PG hydrolases are required to split daughter cells following completion of septum synthesis (Heidrich et al., 2002). A portion of these hydrolases generate PG fragments that are imported into the cell and recycled, a complex pathway that is not essential for growth in laboratory conditions but bears important roles in (i) minimizing energy costs, (ii) sensing the appropriate balance between synthetic and hydrolytic activities, which may be altered by β-lactam antibiotics and other toxic agents, and (iii) avoiding the release of proinflammatory molecules recognized by the host immune system (Bastos et al., 2020; Johnson et al., 2013). PG hydrolases specifically acting on each of the ten amide, ether, and glycosidic bonds present in the PG polymer have been described and most cleavage specificities involve multiple enzymes (supplementary Fig. S6A). Enzymes of different specificities can at least in part compensate for each other, *e.g*. lytic glycosyltransferases and amidases both contribute to the separation of daughter cells (Heidrich et al., 2002; van Heijenoort, 2011). In this study, we show that endopeptidases are specifically required for bacterial growth not only in the context of the formation of 4→3 cross-links by PBPs but also in the context of the formation of 3→3 cross-links by YcbB (Table 3). We also identify for the first time the minimum sets of endopeptidases for each mode of PG cross-linking, namely MepM for 4→3 cross-links and MepM plus MepK for 3→3 cross-links. Endopeptidase overproduction resulting from expression of the genes under the control of heterologous promoters revealed potential functional redundancies in the endopeptidase families. In particular, overproduction of MepH, MepS, or PBP7 compensated for the absence of MepM in the context of a 4→3 cross-linked PG. Overproduction of MepS compensated for the absence of MepM or MepK for growth with a 3→3 cross-linked PG. Overproduction of MepM prevented growth probably due to unbalanced synthesis and hydrolysis of PG cross-links (observed for both 4→3 and 3→3 cross-linked PG). Production of catalytically inactive endopeptidases suggested that MepA and MepS are negatively regulated by Prc-mediated proteolysis, as previously established for MepS (Lai et al., 2017; Singh et al., 2015).

**Table 3.**
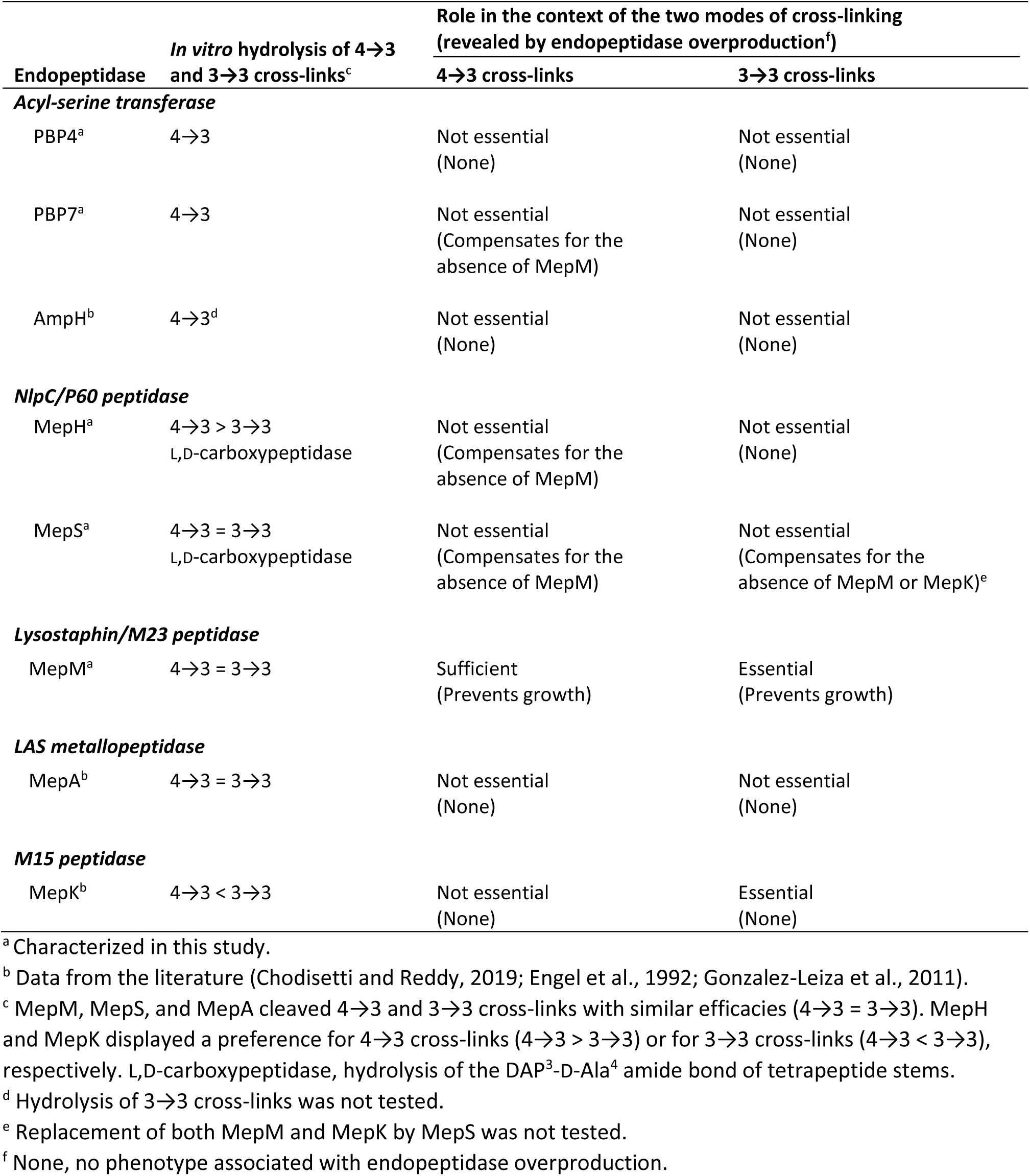
Characteristics of the endopeptidases.

| Endopeptidase | *In vitro* hydrolysis of 4→3 and 3→3 cross-links<sup>c</sup> | Role in the context of the two modes of cross-linking (revealed by endopeptidase overproduction<sup>f</sup>) |  |
| --- | --- | --- | --- |
|  |  | 4→3 cross-links | 3→3 cross-links |
| ***Acyl-serine transferase*** |  |  |  |
| PBP4<sup>a</sup> | 4→3 | Not essential<br>(None) | Not essential<br>(None) |
| PBP7<sup>a</sup> | 4→3 | Not essential<br>(Compensates for the absence of MepM) | Not essential<br>(None) |
| AmpH<sup>b</sup> | 4→3<sup>d</sup> | Not essential<br>(None) | Not essential<br>(None) |
| ***NlpC/P60 peptidase*** |  |  |  |
| MepH<sup>a</sup> | 4→3 > 3→3<br>L,D-carboxypeptidase | Not essential<br>(Compensates for the absence of MepM) | Not essential<br>(None) |
| MepS<sup>a</sup> | 4→3 = 3→3<br>L,D-carboxypeptidase | Not essential<br>(Compensates for the absence of MepM) | Not essential<br>(Compensates for the absence of MepM or MepK)<sup>e</sup> |
| ***Lysostaphin/M23 peptidase*** |  |  |  |
| MepM<sup>a</sup> | 4→3 = 3→3 | Sufficient<br>(Prevents growth) | Essential<br>(Prevents growth) |
| ***LAS metallopeptidase*** |  |  |  |
| MepA<sup>b</sup> | 4→3 = 3→3 | Not essential<br>(None) | Not essential<br>(None) |
| ***M15 peptidase*** |  |  |  |
| MepK<sup>b</sup> | 4→3 < 3→3 | Not essential<br>(None) | Essential<br>(None) |

<sup>a</sup> Characterized in this study.<sup>b</sup> Data from the literature (Chodiseti and Reddy, 2019; Engel et al., 1992; Gonzalez-Leiza et al., 2011).<sup>c</sup> MepM, MepS, and MepA cleaved 4→3 and 3→3 cross-links with similar efficacies (4→3 = 3→3). MepH and MepK displayed a preference for 4→3 cross-links (4→3 > 3→3) or for 3→3 cross-links (4→3 < 3→3), respectively. L,D-carboxypeptidase, hydrolysis of the DAP<sup>3</sup>-D-Ala<sup>4</sup> amide bond of tetrapeptide stems.<sup>d</sup> Hydrolysis of 3→3 cross-links was not tested.<sup>e</sup> Replacement of both MepM and MepK by MepS was not tested.<sup>f</sup> None, no phenotype associated with endopeptidase overproduction.

### Specificity of purified endopeptidases for 4→3 and 3→3 cross-links

The cleavage specificity of the endopeptidases was determined by mass spectrometry (Fig. 5, 7, and S2). Endopeptidases of the PBP family were specific to 4→3 cross-links. Endopeptidases belonging to other families cleaved both 4→3 and 3→3 cross-links with similar efficacies (MepM, MepS, MepA) or with a preference for 4→3 (MepH) or 3→3 (MepK) cross-links (Fig. 5 and 7) (Chodisetti and Reddy, 2019; Engel et al., 1992). This is unexpected since 4→3 and 3→3 cross-links contain amide bonds connecting two R stereo centers (D-Ala^4^→DAP^3^) or an S to an R stereo center (DAP^3^→DAP^3^), respectively (supplementary Fig. S6A). This could imply that endopeptidases of the Mep families mainly interact with the acceptor stems of cross-linked muropeptides, which are the same for both types of cross-links, whereas endopeptidases of the

PBP family interact with a donor tetrapeptide stem only present in 4→3 cross-linked muropeptides (supplementary Fig. S6B).

### Integration of endopeptidases into the global regulation of 4→3 and 3→3 PG cross-linking

D,D-carboxypeptidases, which cleave off the terminal residue (D-Ala^5^) of pentapeptide stems, are thought to negatively control the transpeptidase activity of PBPs since these enzymes require a pentapeptide donor (Fig. 1). A less studied impact of D,D-carboxypeptidases is the formation of the essential tetrapeptide donor substrate of the LDTs, except for two publications reporting that PBP5 and PBP6a are essential for YcbB-mediated β-lactam resistance and for rescue of a defect in lipopolysaccharide synthesis, respectively (Hugonnet et al., 2016; Morè et al., 2019). Thus, D,D-carboxypeptidases have crucial roles in controlling the relative contributions of transpeptidases of the D,D and L,D specificities to PG cross-linking by both decreasing access of PBPs to pentapeptide stems and increasing access of LDTs to tetrapeptide stems.

Endopeptidases participate in the metabolism of PG cross-links in several ways. MepH and MepS display L,D-carboxypeptidase activity leaving tripeptides as the main (> 80%) end product of *in vitro* PG hydrolysis (Fig. 7). This activity may negatively control the L,D-transpeptidase activity of YcbB by hydrolysis of D-Ala^4^ thereby preventing access to its tetrapeptide donor. In addition, hydrolysis of 4→3 cross-linked Tetra→Tetra and Tetra→Tri dimers by the endopeptidases generates free tetrapeptide stems (Fig. 5 and 7). These tetrapeptide stems can be used as donor by YcbB for formation of 3→3 cross-links, as demonstrated for the LDTs of *Mycobacterium smegmatis* (Baranowski et al., 2018). In contrast, the D,D-transpeptidase activity of PBPs exclusively relies on *de novo* synthesis and translocation of pentapeptide-containing precursors since the D-Ala^5^ residue of pentapeptide stems is rapidly cleaved off by D,D-carboxypeptidases if they are not used for formation of 4→3 cross-links. Thus, YcbB is expected to function as a rescue enzyme to restore cross-linking in regions of the PG that are compromised by 4→3 or 3→3 endopeptidases. This mechanism was proposed for PG reparation following disassembly of the lipopolysaccharide export machinery that crosses the PG layer (Morè et al., 2019). The combined activities of endopeptidases cleaving 4→3 cross-links and of L,D-transpeptidases could contribute to the enrichment in 3→3 cross-links in stationary phase cultures (Pisabarro et al., 1985). In turn, this enrichment may protect cells from hydrolases active on 4→3 cross-linked PG. Previous analyses proposed that two L,D-transpeptidases may contribute to the enrichment of PG in 3→3 cross-links, namely YcbB (LdtD), induced by the cell envelope Cpx stress system, and YnhG (LdtE), expressed under the control of sigma S and induced in stationary phase (Delhaye et al., 2016; Weber et al., 2005).

### Participation of YcbB to PG polymerization complexes

PG polymerization is generally thought to be performed by two multiprotein complexes involved in the expansion of the lateral cell wall (elongasome) and in the formation of the septum (divisome) (Pazos et al., 2017). Replacement of the D,D-transpeptidase activity of all PBPs by the L,D-transpeptidase activity of YcbB raises several questions regarding the identity of the partners of YcbB for the assembly of lateral wall and septum PG, and whether YcbB physically replaces PBPs in the PG polymerization complexes. Our data support a model in which YcbB functions with two different sets of partners for lateral wall and septum PG assembly as follows.

For the assembly of lateral wall PG, inactivation of the transpeptidase domain of PBP2 by β-lactams leads to uncoupling of the transglycosylation and transpeptidation reactions, the former being most probably catalyzed by RodA (Cho et al., 2014; Uehara and Park, 2008). Uncross-linked glycan chains accumulate in the periplasm and are eventually cleaved by the Slt70 lytic transglycosylase and recycled. According to the model presented in Fig. 12, and in agreement with a previous study (Cho et al., 2014), the reactions catalyzed by Slt70 and YcbB occur in competition implying that YcbB-mediated cross-linking is not coupled to glycan chain polymerization by RodA. This also implies that YcbB could function in the PG layer in combination with the MepM endopeptidase known to participate in cell elongation (Banzhaf et al., 2020; Singh et al., 2012; Truong et al., 2020; Uehara et al., 2009). In agreement with this model, impaired Slt70 activity had a positive impact on β-lactam resistance mediated by YcbB. This was established both by the selection of mutations enabling expression of β-lactam resistance in liquid medium, which mapped in the *sltY* gene encoding Slt70 (Table 2) and by the deletion of *sltY*, which lowered the level of *ycbB* expression required for resistance (Fig. 11).

**Figure 12.**
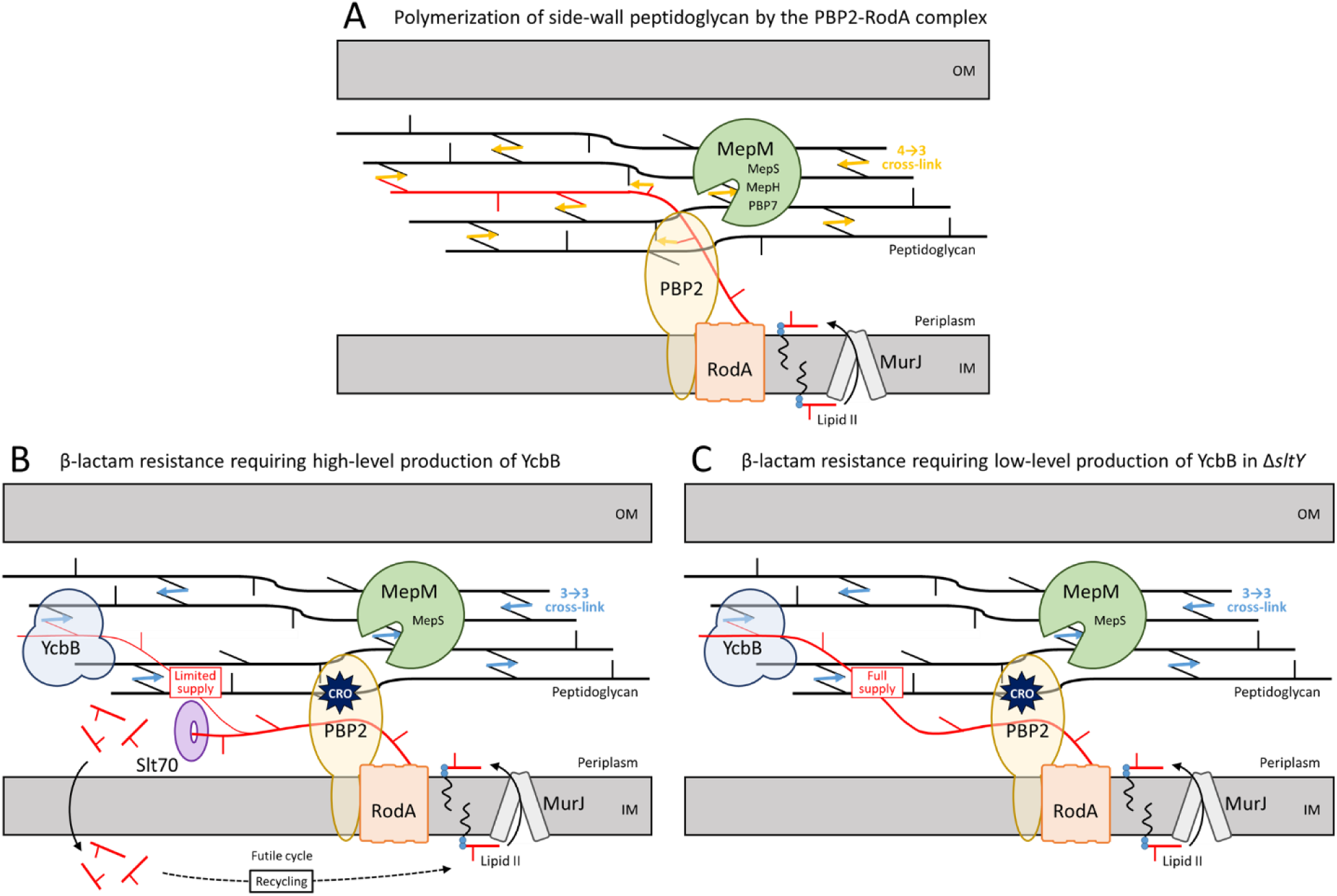
Proposed model for polymerization of side wall PG by transpeptidases of the D,D- or L,D-specificity. (**A**) In wild-type cells, the disaccharide pentapeptide subunit linked to the undecaprenyl lipid transporter (Lipid II) is translocated to the outer leaflet of the cytoplasmic membrane by MurJ and polymerized by the glycosyltransferase and D,D-transpeptidase activities of the PBP2-RodA complex. MepM is essential and sufficient for insertion of new material in the PG net, although this endopeptidase can be replaced by overproduction of MepH, MepS, or PBP7. (**B**) Inhibition of PBP2 by β-lactams leads to the accumulation of uncross-linked glycan chains that are cleaved by the Slt70 lytic transglycosylase. This limits the supply of glycan chains for cross-linking by the YcbB L,D-transpeptidase. Under this condition, MepM or overproduction of MepS is required for insertion of new glycan strands. (**C**) Deletion of the *sltY* gene encoding lytic transglycosylase Slt70 prevents digestion of uncross-linked glycan chains leading to a full supply of neo-synthesized glycan chains to YcbB and improved expression of β-lactam resistance. IM, inner membrane; OM, outer membrane; CRO, ceftriaxone.

For the assembly of septum PG, YcbB was proposed to cooperate with the glycosyltransferase activity of Class A PBP1b (Caveney et al., 2019; Hugonnet et al., 2016). In support of this hypothesis, microscale thermophoresis experiment revealed that purified YcbB interacts with PBP1b and PBP5 (D,D-carboxypeptidase). Furthermore, the glycosyltransferase activity of PBP1b is essential for YcbB-mediated β-lactam resistance whereas the combined deletion of class A PBP1a and PBP1c had no impact. We cannot rule out the possibility that the glycosyltransferase activity of FtsW also contributes to septum PG polymerization in the presence of β-lactams but this would require that the inactivation of PBP3 by β-lactams lead to the uncoupling of glycan chain polymerization and PG cross-linking, as proposed for the RodA-PBP2 complex (see above). This is not supported by the analyses of PG recycling in conditions of selective inhibition of PBP3 by aztreonam (Uehara and Park, 2008). The endopeptidases involved in septum formation have not been identified, except for a contribution of PBP4, which has an effect on the timing of septation (Verheul et al., 2020). MepK is a candidate for this function in 3→3 cross-linked PG although this is currently not supported by any experimental evidence and it remains to be seen if endopeptidases are needed for septum PG synthesis.

**Figure S6.**
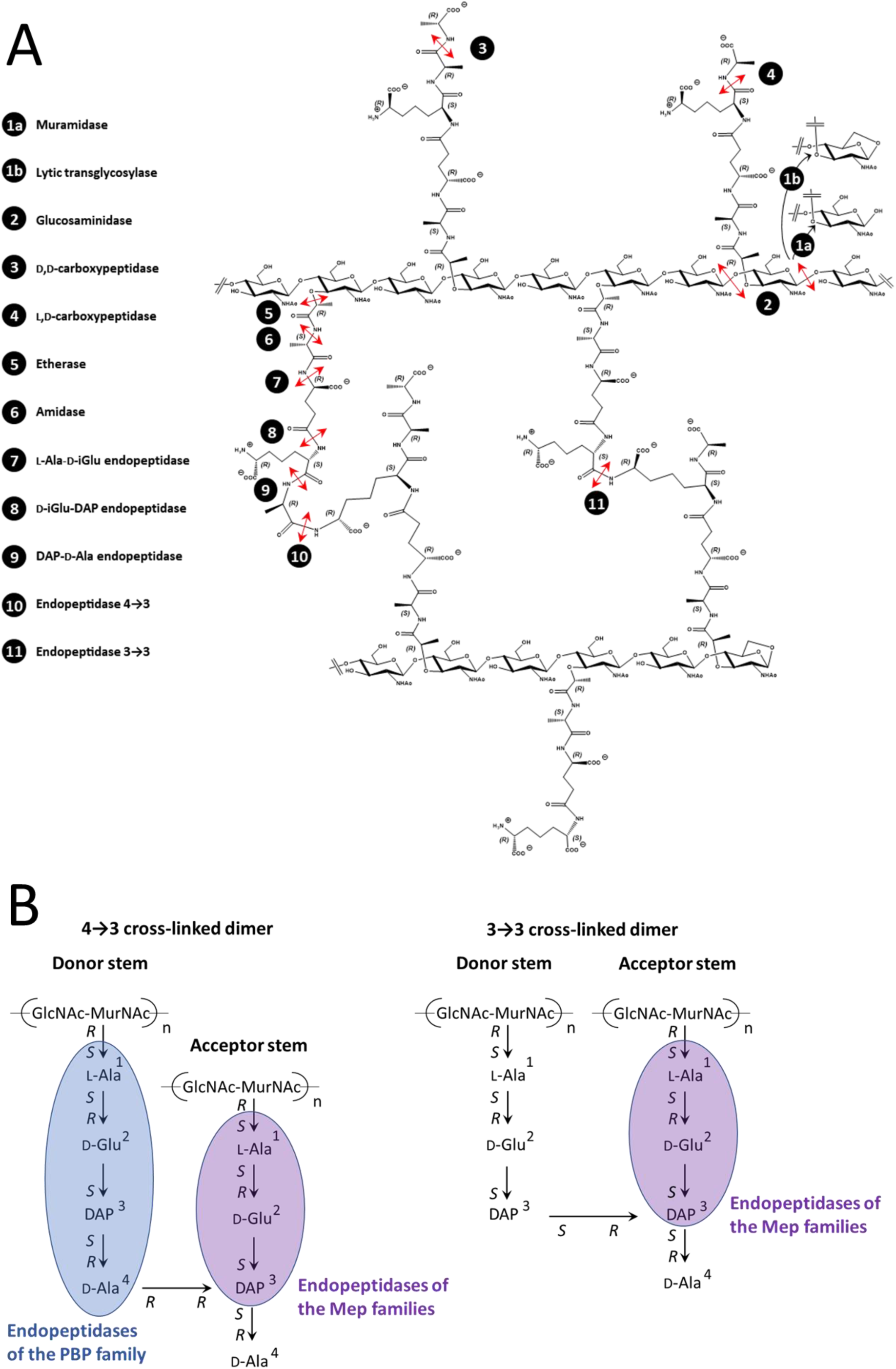
Specificity of PG hydrolases. (**A**) Highlight of enzyme stereospecificity. The commonly used endopeptidase designation was employed in the entire manuscript to refer to the cleavage of internal bonds although certain enzymes do not cleave peptide bonds connecting the α amino and carboxyl groups of two consecutive amino acids and should have been more precisely referred to as amidases. (**B**). Recognition of the donor and acceptor stems of dimers by PBP and Mep endopeptidases accounting for the 4→3 *versus* 4→3 plus 3→3 specificities. According to this model, endopeptidases of the Mep families cleave both 4→3 plus 3→3 cross-links since they interact with the tripeptide portion of the acceptor stem, which is present in both types of dimers. In contrast, endopeptidases of the PBP families specifically interact with the tetrapeptide donor stem of 4→3 cross-linked dimers.

## MATERIALS AND METHODS

### Strains, plasmids, and growth conditions

All strains were derived from *E. coli* BW25113 (Baba et al., 2006). The origin and characteristics of plasmids are listed in Supplementary Table S1. Bacteria were grown in brain heart infusion (BHI; Difco) broth or agar at 37 °C unless otherwise specified. Liquid cultures were performed with aeration (180 rpm). The growth media were systemically supplemented with drugs to counter-select plasmid loss (Supplementary Table S1). The same drugs at the same concentrations were used to select transformants. Kanamycin at 50 μg/ml was used for the Km^R^ cassette. Induction of the *lacZYA, araBAD*, and *rhaBAD* promoters was performed with isopropyl β-D-1-thiogalactopyranoside (IPTG, 40 or 50 μM), L-arabinose (0.2 or 1%), and L-rhamnose (0.2 or 1%), respectively. Plasmids constructed in this study were obtained by using NEBuilder HiFi DNA assembly (New England Biolabs) method, unless otherwise specified.

Growth curves were obtained in a 96-well plate using an Infinite 200 PRO microplate reader (TECAN). Briefly, bacteria were grown to the late exponential phase, *i.e*. to an optical density at 600 nm (OD600) greater than 1.0 *(ca*. 6 h at 37 °C under agitation). The OD600 was adjusted to 1.0 and 5 μl were inoculated in 195 μl of BHI broth supplemented with drugs and inducers, as specified in the legends to figures. Growth was monitored at 600 nm every 5 min for 12 h at 37 °C with vigorous shaking.

### Construction of *E. coli* strains carrying gene deletions

The Keio collection comprises 3,985 mutants obtained by replacement of non-essential genes by a kanamycin resistance (Km^R^) cassette (Baba et al., 2006). P1 transduction of the Km^R^ cassette from selected mutants was used to introduce deletions of specific genes involved in PG synthesis (Datsenko and Wanner, 2000). For multiple gene deletions, the Km^R^ cassette was removed by the FLT recombinase encoded by plasmid pCP20. The presence of the expected deletions was confirmed by PCR amplification at each deletion step. Supplementary Fig. S4 shows lineages that have been obtained by parallel serial deletions.

To study the 3→3 mode of cross-linking, plasmid pKT2(*ycbB*) and pKT8(*relA’*) were introduced into the derivatives of *E. coli* BW25113 *ΔrelA* obtained by deletion of various combinations of endopeptidase genes. For the sake of simplicity, the latter strains were referred to as BW25113(*ycbB*, *relA’*) derivatives even though gene deletions preceded the introduction of pKT2(*ycbB*) and pKT8(*relA’*).

### Construction of *E. coli* BW25113 *ΔrelA* Δ8EDs

The plasmid *pHV53(P_rhaBAD_-TIS2-mepM)* was introduced into BW25113 *ΔrelA* Δ7EDs harboring *mepM* as the only chromosomal endopeptidase-encoding gene. The *mepM* deletion was introduced into BW25113 *ΔrelA* Δ7EDs *pHV53(P_rhaBAD_-TIS2-mepM)* in the presence of 0.2% L-rhamnose by P1 transduction as described above. Growth of the resulting BW25113 *ΔrelA* Δ8EDs pHV53(P*rhaBAD-TIS2-mepM)* strain was dependent on the induction of the plasmid copy of *mepM* mediated by L-rhamnose.

### Mutant selection and whole-genome sequencing

*E. coli* BW25113 M1 was streaked for isolated colonies on agar plates containing 10 μg/ml tetracycline to counter-select loss of plasmid pJEH12(*ycbB*). The selection procedure was independently carried out starting with four independent colonies. Briefly, 5 ml of BHI broth supplemented with 10 μg/ml tetracycline and 50 μM IPTG were inoculated with a colony. Bacteria were grown overnight with shaking (180 rpm). A fraction of 1 ml of the culture was inoculated in 250 ml of BHI broth supplemented with 16 μg/ml ampicillin and 50 μM IPTG. Bacteria were grown overnight with shaking and streaked on BHI agar containing 16 μg/ml ampicillin and 50 μM IPTG. A colony was inoculated in 250 ml of BHI broth supplemented with 16 μg/ml ampicillin and 50 μM IPTG. Bacteria were grown overnight with shaking and streaked on BHI agar containing 16 μg/ml ampicillin and 50 μM IPTG. Five ml of BHI broth containing 10 μg/ml tetracycline was inoculated with a single colony and genomic DNA was extracted (Wizard DNA extraction kit, Promega). Genomic DNA was sequenced by paired-end joining Illumina (Biomics Platform of the Institut Pasteur, Paris, France). Identification of the mutations was performed with the *breseq* pipeline (Deatherage and Barrick, 2014).

### Plating efficiency assay

Bacteria were grown to the late exponential phase, *i.e*. to an optical density at 600 nm (OD600) greater than 1.0 *(ca*. 6 h at 37 °C under agitation). The OD600 was adjusted to 1.0 and 10-fold dilutions (10^-1^ to 10^-6^) were prepared in BHI broth. Ten μl of the resulting bacterial suspensions were spotted on BHI agar supplemented with inducers and drugs as indicated in the legend to figures. For the disk diffusion assay, 5 μl of the bacterial suspension adjusted to an OD600 of 1.0 were inoculated in 5 ml of water. BHI agar plates were flooded with the latter suspension, excess liquid was removed, and the plates were kept at room temperature for 15 min prior to the addition of paper disks containing antibiotics or inducers. Plates were imaged after 16 h (or 24 h for plates containing ceftriaxone) of incubation at 37 °C.

### Purification of endopeptidases

The *mepM* gene was amplified by PCR and cloned into pET-TEV between the NdeI and XhoI restriction sites. The fusion protein comprised a 6 x histidine tag, a TEV protease cleavage site, and residues 41-440 of MepM. The *mepH* and *mepS* genes were independently amplified by PCR and cloned in frame with *dsbC* into pETMM82 using NEBuilder HiFi DNA assembly (New England Biolabs). The fusion proteins comprised the DsbC chaperone (Firczuk and Bochtler, 2007), a 6 x histidine tag, a TEV protease cleavage site, and residues 28-271 of MepH or 25-188 of MepS. The enzymes were produced in *E. coli* BL21(DE3) following induction by 0.5 mM IPTG for 18 h at 16 °C. The endopeptidases were purified in 50 mM Tris-HCl pH 8.0 from a clarified lysate by nickel affinity chromatography (elution with 0.5 mM imidazole). The endopeptidases were dialyzed overnight at 4 °C against 50 mM Tris-HCl pH 8.0, 0.5 mM EDTA. N-terminal tags were cleaved overnight at room temperature following addition of 10 μg of TEV protease for every mg of protein and DTT at a final concentration of 0.5 mM. MepM, MepH, and MepS were further purified by size-exclusion chromatography (Superdex 75 HiLoad 26/60, GE Healthcare) in 50 mM Tris-HCl pH 7.5, 200 mM NaCl.

PBP4 was purified from strain BL21(DE3) pET21bΩPBP4Δ1-60 as previously reported (Banzhaf et al., 2020). Briefly, cells were grown in the presence of 1 mM IPTG for 8 h at 20 °C and then harvested by centrifugation at 7,500 × *g*, 4 °C, 15 min. Cell pellets were resuspended in 50 mM Tris-HCl pH 8.0, 300 mM NaCl, and lysed by sonication. Following centrifugation at 14,000 × *g*, 1 h, 4 °C, the NaCl concentration was reduced by stepwise dialysis in a Spectra/Por dialysis membrane (MWCO 12-14 kDa) against 50 mM Tris-HCl pH 8.5 containing (i) 200 mM NaCl, (ii) 100 mM NaCl, and (iii) 30 mM NaCl. The sample was centrifuged at 7,500 × *g*, 4 °C, 10 min and the supernatant applied to a 5 ml HiTrap Q HP IEX column in 25 mM Tris-HCl pH 8.5, 30 mM NaCl. Protein was eluted from the column with a linear gradient from 50 mM Tris-HCl pH 8.5, 100 mM NaCl, to 25 mM Tris-HCl pH 8.0, 1 M NaCl, over a 100 ml volume. Fractions containing PBP4 were combined and dialyzed against 10 mM potassium phosphate pH 6.8, 300 mM NaCl. Protein was applied at 1 ml/min to a 5 ml ceramic hydroxyapatite column (BioRad BioscaleTM) in the dialysis buffer, and a 50 ml linear gradient to 500 mM potassium phosphate pH 6.8, 300 mM NaCl, was applied. Fractions with PBP4 were dialyzed overnight against 25 mM HEPES-NaOH pH 7.5, 300 mM NaCl, 10% glycerol, and concentrated to *ca*. 5 ml using a Vivaspin concentrator spin column (Sartorius). The protein sample was applied to a HiLoad 16/600 Superdex 200 column (GE healthcare) at 1 ml/min and eluted in a linear gradient to 25 mM HEPES-NaOH pH 7.5, 300 mM NaCl, 10% glycerol. The collected fractions containing PBP4 were combined.

PBP7 was purified from strain BL21(DE3) pET28aΩ*pbpG*Δ1-75 as previously reported (Banzhaf et al., 2020). Briefly, cells were grown in the presence of 1 mM IPTG for 3 h at 30 °C before being harvested by centrifugation and resuspended in 25 mM Tris-HCl pH 7.5, 500 mM NaCl, 20 mM imidazole. Following sonication and subsequent centrifugation, the lysate was applied to a 5 ml HisTrap HP column (GE healthcare) and washed with 4 column volumes of 25 mM Tris-HCl pH 7.5, 500 mM NaCl, 20 mM imidazole. Bound protein was eluted with 25 mM Tris-HCl pH 7.5, 300 mM NaCl, 400 mM Imidazole. Elution fractions containing PBP7 were combined and dialyzed overnight against 25 mM HEPES-NaOH pH 7.5, 300 mM NaCl, 10% glycerol, in the presence of 1 unit/ml of restriction grade thrombin (Novagen) to remove the oligohistidine tag. The sample was then concentrated to *ca*. 5 ml using a Vivaspin concentrator spin column (Sartorius) at 4,500 × *g*, 4 °C. The protein sample was applied to a HiLoad 16/600 Superdex 200 column (GE healthcare) at 1 ml/min and eluted in 25 mM HEPES-NaOH pH 7.5, 300 mM NaCl, 10% glycerol. Elution fractions containing PBP7 were combined.

Protein concentrations were determined by the Bio-Rad protein assay using bovine serum albumin as a standard. Endopeptidases were stored at −80 °C.

### Preparation of sacculi

Bacteria were grown in M9 minimal medium supplemented with 0.1% glucose at 37 °C for 48 h. Bacteria were harvested by centrifugation and boiled in 4% sodium dodecyl sulfate (SDS) for 1 h. Sacculi were harvested by centrifugation (20,000 x *g* for 20 min at 20 °C), washed five times with water, and incubated with 100 μg/ml pronase overnight at 37 °C in 20 mM Tris-HCl pH 7.5. Sacculi were washed five times with water and incubated overnight at 37 °C with 100 μg/ml trypsin in 20 mM sodium phosphate pH 8.0. Sacculi were washed five times with water, boiled for 5 min, collected by centrifugation, resuspended in water, and stored at −20 °C.

### Digestion of sacculi

Sacculi were digested overnight at 37 °C with 120 μM lysozyme alone or in association with an endopeptidase in 40 mM Tris-HCl pH 8.0. Insoluble material was removed by centrifugation and the soluble fraction containing muropeptides was reduced with sodium borohydride for 1 h in 125 mM borate buffer pH 9.0. The pH of the solution containing the reduced muropeptides was adjusted to 4.0 with phosphoric acid. Muropeptides were separated by *rp*HPLC in a C18 column (Hypersil GOLD aQ; 250 x 4.6 mm; 3 μm, Thermoscientific) at a flow rate of 1 ml/min with a linear gradient (0 to 20%) applied between 10 and 60 min (buffer A, TFA 0.1%; buffer B, acetonitrile 20% TFA 0.1%). Absorbance was monitored at 205 nm and fractions were collected, lyophilized, and analyzed by mass spectrometry. Mass spectra were obtained on a Bruker Daltonics maXis high-resolution mass spectrometer (Bremen, Germany) operating in the positive mode (Analytical Platform of the Muséum National d’Histoire Naturelle, Paris, France).

**Table S1.**
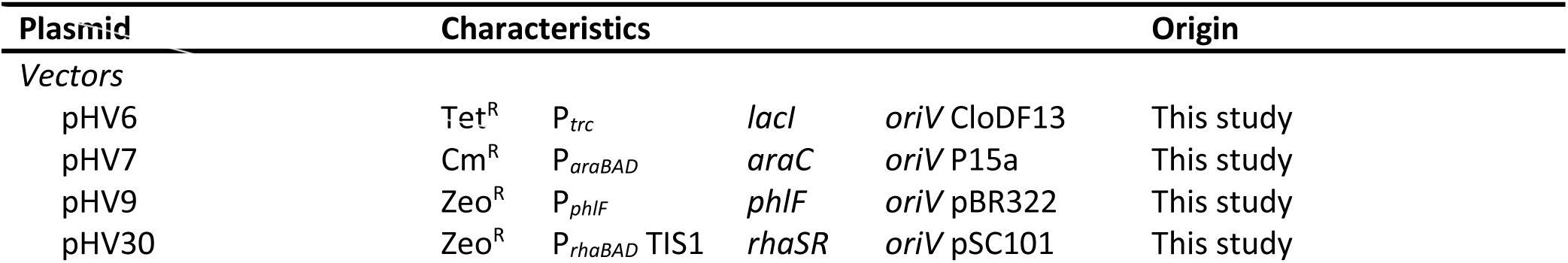
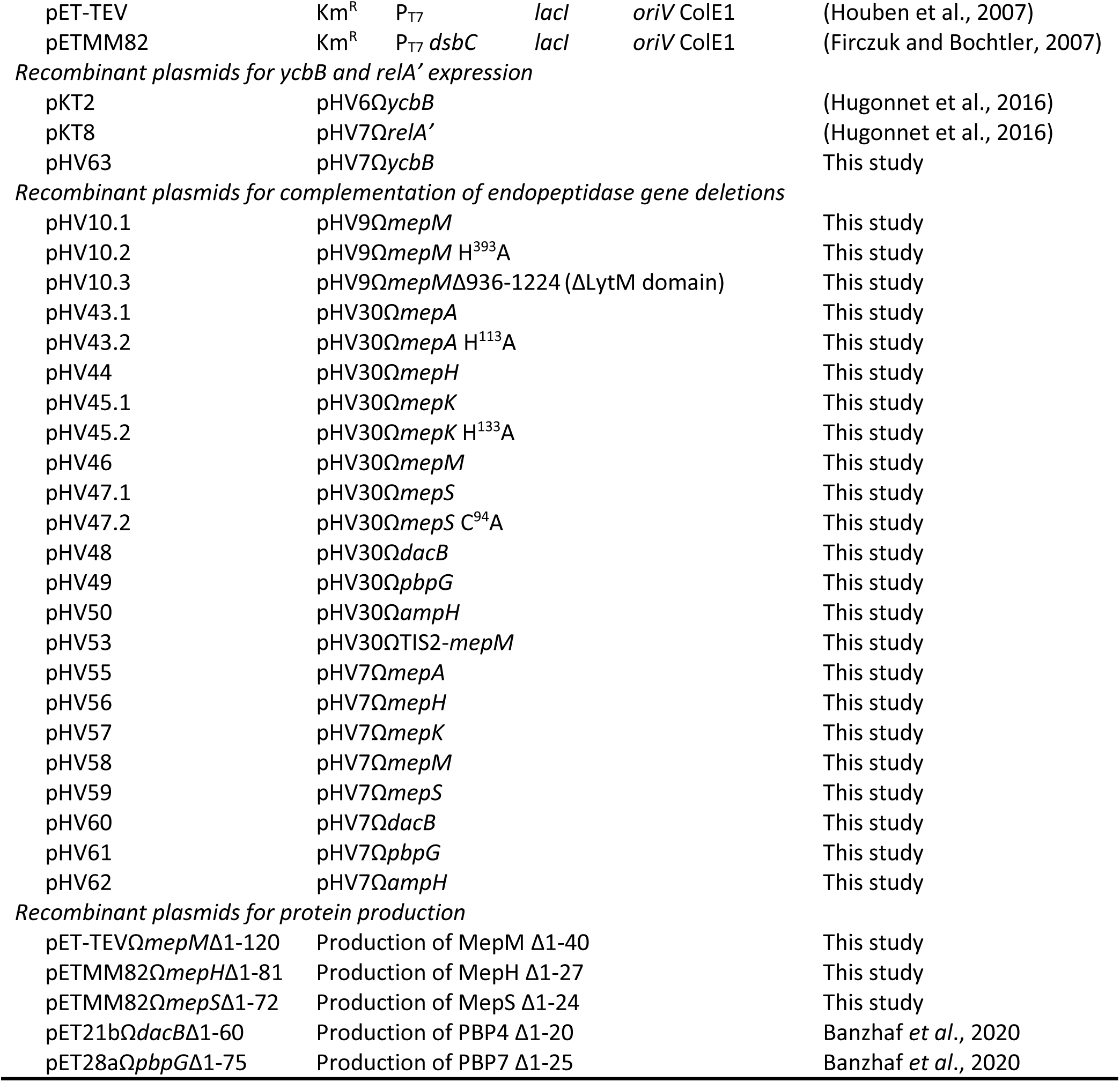
Characteristics and origin of the plasmids used in this study.

**Table S2.** Characteristics and origin of *E. coli* strains used in this study.

| Strain | Characteristics | Origin |
| --- | --- | --- |
| BW25113 | Δ(*araD-araB*)567 Δ(*rhaD-rhaB*)568<br>Δ*lacZ*4787 (::rrnB-3) *hsdR*514 *rph*-1 | (Baba et al., 2006) |
| BW25113(*ycbB*, *relA'*) | Δ*relA* derivative of BW25113 harboring pKT2(*ycbB*)<br>and pKT8(*relA'*) | (Hugonnet et al., 2016) |
| BW25113 M1 | Ampicillin resistant derivative of BW25113<br>pJEH12(*ycbB*) harboring a mutation in the 5' UTR<br>region of *ileRS* | (Hugonnet et al., 2016) |
| BL21(DE3) | Host for protein production | (Wood, 1966) |

## ACKNOWLEDGEMENTS

This work was supported by the French National Research Agency ANR ‘RegOPeps’ (grant ANR-19-CE44-0007 to JEH). We thank L. Dubost and A. Marie for technical assistance in the collection of mass spectra. We also thank L. Ma and R. Legendre for technical assistance in genome sequencing. The Biomics Platform is a member of the ‘France Génomique’ consortium supported by the French National Research Agency ANR (grant ANR-10-INBS-0009) and IBISA. We also thank Z. Edoo for proofreading the manuscript.

